# Epigenomic signature of the progeroid Cockayne syndrome exposes distinct and common features with physiological ageing

**DOI:** 10.1101/2021.05.23.445308

**Authors:** Clément Crochemore, Claudia Chica, Paolo Garagnani, Giovanna Lattanzi, Steve Horvath, Alain Sarasin, Claudio Franceschi, Maria Giulia Bacalini, Miria Ricchetti

## Abstract

Cockayne syndrome (CS) and UV-sensitivity syndrome (UVSS) are rare genetic disorders caused by mutation of the DNA repair and chromatin remodelling proteins CSA or CSB, but only CS patients display a progeroid and neurodegenerative phenotype. As epigenetic modifications constitute a well-established hallmark of ageing, we characterized genome-wide DNA methylation (DNAm) of fibroblasts from CS *versus* UVSS patients and healthy donors. The analysis of differentially methylated positions and regions revealed a CS-specific epigenetic signature, enriched in developmental transcription factors, transmembrane transporters, and cell adhesion factors. The CS-specific signature compared to DNAm changes in other progeroid diseases and regular ageing, identifyied commonalities and differences in epigenetic remodelling. CS shares DNAm changes with normal ageing more than other progeroid diseases do, and according to the methylation clock CS samples show up to 13-fold accelerated ageing. Thus, CS is characterized by a specific epigenomic signature that partially overlaps with and exacerbates DNAm changes occurring in physiological aging. Our results unveil new genes and pathways that are potentially relevant for the progeroid/degenerative CS phenotype.

## Introduction

In several rare genetic disorders ageing is greatly accelerated^1^, but the molecular defects leading to precocious ageing are still unknown. Progeroid syndromes are genetically, biochemically, and clinically heterogeneous. While Hutchinson–Gilford progeria syndrome (HGPS) is due to a defective nuclear lamina as a consequence of mutations in *LMNA* gene, Cockayne syndrome (CS) and Werner syndrome (WS) result from mutation in multifunctional proteins best known for their role in DNA repair. It is not yet known which genes on chromosome 21 contribute to the progeroid phenotype of trisomy 21 also known as Down syndrome (DS). The extent and combination of ageing-related defects^2^ are specific to each progeroid syndrome. Since not all alterations observed during regular ageing are clinically recapitulated, these conditions are often referred to as *segmental* progeroid syndromes^3^.

Here we focus on CS which is a devastating multisystemic disease characterized by neurodevelopmental defects, precocious ageing, and in most cases cutaneous photosensitivity. It is a progressive disorder with most symptoms appearing relatively early in life and worsening with time^4^. Multiple degrees of severity have been described, including a prenatal-onset form known as cerebro-oculo-facioskeletal (COFS) syndrome, an early-onset severe form (CS type II, average age of death 5-6 years), a classical or moderate form (CS type I, mean life expectancy around 16 years) and a mild or late-onset form (CS type III)^5^.

CS is due to mutations in either *CSB* or *CSA* genes (approximately 70:30 ratio)^5,6^, which code repair factors of UV-induced DNA damage through the transcription-coupled nucleotide excision repair (TC-NER) pathway^7^. CS defects, including the precocious ageing phenotype, have been essentially ascribed to impaired DNA repair. Nevertheless, mutations in *CSB* or *CSA* are also responsible for an independent syndrome, the UV-sensitive syndrome (UVSS) that is characterized by severe photosensitivity but no sign of premature ageing^8,9^. About 8 cases of UVSS, which display a similar phenotype, are known worldwide and some of which are due to mutations in UVSSA, a protein that acts in TC-NER in concert with CSA and CSB^10,11^. These case reports *de facto* uncouple the DNA repair defect from precocious ageing and suggest that impairment of other functions of multifunctional CSA and CSB proteins are responsible for progeroid defects.

Indeed, CSA and CSB are also transcription factors^12^, are implicated in chromatin remodelling^13,14^, and have been detected in mitochondria where they act in repairing oxidative damage^15^ and promoting transcription^16^ of mitochondrial DNA. Additionally, we showed that in CS (but not UVSS) cells the CSA/CSB impairment results in overexpression of the HTRA3 protease leading to degradation of the mitochondrial DNA polymerase POLG1, and consequent mitochondrial dysfunction^17^. CSA and CSB mutations represent therefore an exceptional case for disentangling downstream genes and pathways involved in dramatically different phenotypes, since their impairment results (CS) or not (UVSS) in accelerated ageing. We recently associated the CS-specific HTRA3/POLG1/mitochondrial alterations with replicative senescence in normal cells, and identified CSB depletion as an early trigger of this process, mechanistically linking progeroid defects with a major ageing-related process^18^.

Different molecular hallmarks of ageing (*e.g*. epigenetic alterations, senescence, and mitochondrial dysfunction) have been identified^19^. DNA methylation, an epigenetic mechanism corresponding to the addition of a methyl group in the fifth position of a cytosine could contribute to the ageing process since it is associated with important changes in gene expression^20^. DNA methylation globally decreases with ageing, largely as a consequence of demethylation of repetitive retroelements (LINEs, Alu sequences), driving genomic instability^21^. In recent years, epigenome-wide association studies (EWAS) performed in ageing contexts demonstrated that hypo-/hyper-methylation of specific sets of CpGs can be used to build epigenetic clocks, which are predictive of chronological age or mortality risk (*i.e*, the phenotypic age or GrimAge of an individual)^22–24^.

Important alterations of the DNA methylation profile were described in progeroid WS-, HGPS-, and DS-derived cells compared to healthy controls^25–28^, and were associated with increased epigenetic age^25,29,30^. We performed genome-wide analysis of the methylome of CS and UVSS patients, with the aim of discriminating CS-specific ageing-related DNA methylation changes by filtering out the UVSS condition (mutated but non-progeroid). This analysis identified genes enriched in three major categories, many of which have not been previously associated with ageing. We also demonstrated a dramatic increase of the epigenetic age of CS fibroblasts, and comparison with available datasets revealed a CS epigenetic signature closer to physiological ageing than HGPS, WS or DS, underscoring CS as a relevant model of accelerated ageing.

## Results

### Genome-wide DNA methylation of fibroblasts from CS, UVSS and healthy subjects

We analysed genome-wide DNA methylation in dermal fibroblasts isolated from 7 CS patients, 2 UVSS patients and 3 healthy subjects, with no siblings, and at similar passage number. An overview of the study design is reported in Fig S1. Samples were processed using the Infinium HumanMethylation450 BeadChip or Infinium MethylationEPIC BeadChip microarrays, as indicated in Figure 1A, and seven samples were assessed using both platforms. After data pre-processing (see Materials and Methods), we selected the probes that were in common between the two arrays (452567 probes). Principal component analyses (PCA) showed a separation of samples according to the type of platform on which they had been processed (Fig. S2A); within each batch, samples tended to cluster according to the pathological state (Fig. S2B). After applying the *ComBat* algorithm to adjust for the platform effect, no differences between the 450k and EPIC samples were observed (Fig. S2C), and samples still clustered together according to the pathological state (Fig. S2D). Indeed, along the first component UVSS cells were closer to WT than CS-II (severe form of CS) and, to a larger extent, CS-I cells (mild form of CS). Moreover, the second component identified a clear separation between CS-I cells mutated for *CSA* and *CSB*, which, to date, are clinically indistinguishable^5^. However, CS-I CSA and CS-I CSB derived from male and female patients, respectively. We therefore removed the probes mapping on X and Y chromosomes (10585 probes) and performed again the PCA analysis, which returned a result substantially overlapping the previous one (Fig. S2E-H). Distinct CS-I CSA and CS-I CSB clusters were confirmed also upon removal of 1184 autosomic probes whose methylation level has been shown to be affected by sex in whole blood^31^ (Fig. S2I-L). The global pattern of methylation values was not affected by the ComBat batch correction (Fig. S2M and N).

**Figure 1.**
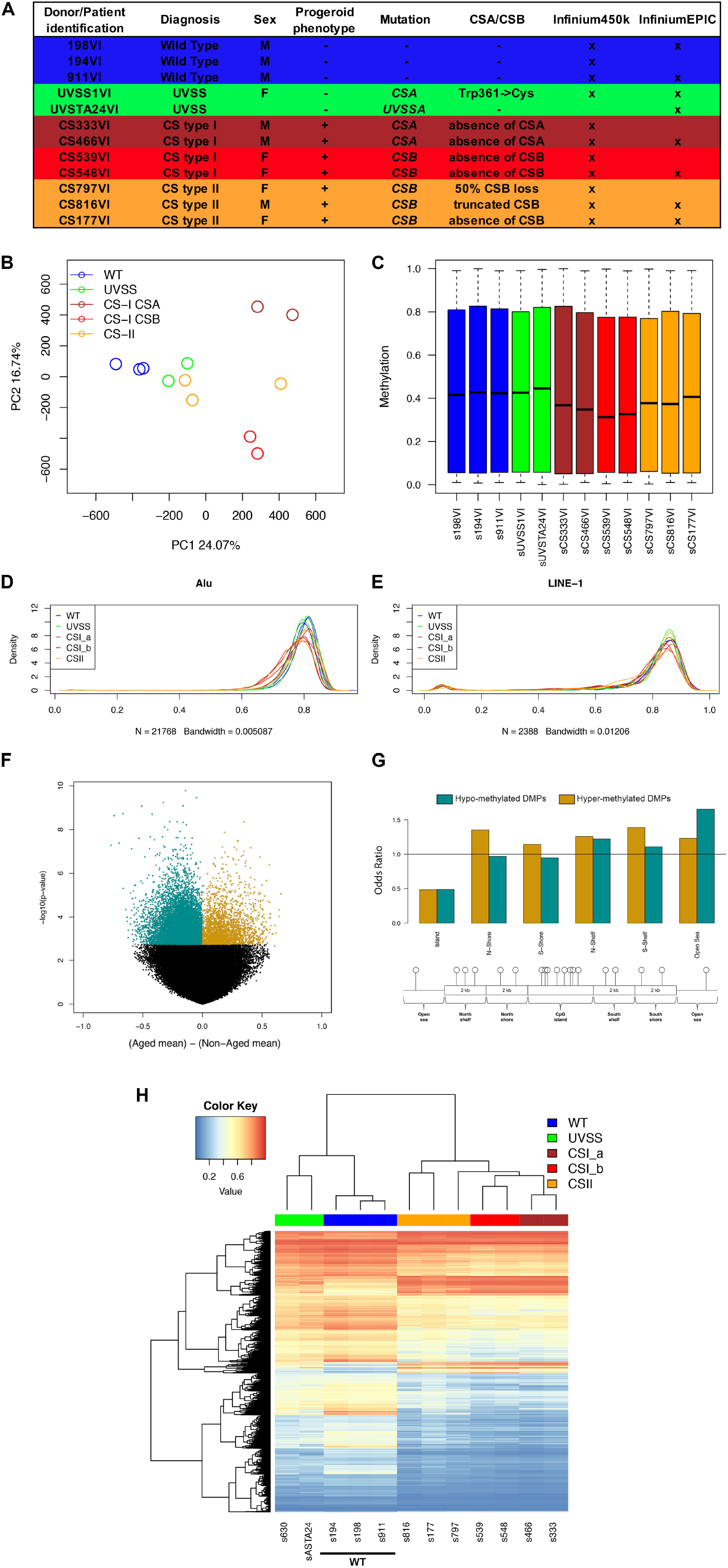
Global features of DNA methylation in CS skin fibroblasts. (**A**) Characteristics of primary skin fibroblasts derived from three healthy donors (WT), two patients with UVSS and seven patients with CS. A cross the platform used for the DNA methylation analysis (with 7 samples analysed with both platforms). (**B**) Principal Component Analysis (PCA) of DNA methylation levels of probes assessed in the Infinium450k and InfiniumEPIC beadchips (merged dataset), after batch correction; dots are coloured according to the pathological or control condition: WT, UVSS, and CS. (**C**) Boxplot of global DNA methylation values in WT, UVSS, and CS cells in the merged dataset after batch correction. Density of the methylation beta values of each sample for the repetitive Alu (**D**) and LINE-1 (**E**) elements. (**F**) Volcano plot showing nominal p-values *vs* the difference in mean methylation values between the Aged and the Non-Aged group. Green: hypomethylation; gold: hypermethylation (**G**) Enrichment of DMPs in different genomic regions. Odds ratio values are reported on the y-axis. Green: hypomethylation; gold: hypermethylation (**H**) Heatmap and hierarchical clustering on DMPs methylation values in the various patient and donor-derived cells.

We performed the subsequent analyses using the batch-corrected data, removing the probes on X and Y chromosomes (441982 probes left) and, for the 7 replicated samples, averaging methylation values from the 450k and EPIC platforms. PCA and boxplots calculated on this merged dataset are reported in Figure 1B and Figure 1C, respectively. Interestingly, boxplots of the methylation values showed a global hypomethylation in CS cells compared to WT and UVSS (Fig. 1C). Moreover, analysis of the DNA methylation of repetitive sequences unveiled an important hypomethylation of Alu sequences in CS samples, and in particular in CS-I, compared to WT and UVSS (Fig. 1D). Conversely the DNA methylation pattern of LINE-1 sequences did not appear to be correlated with the pathological state (Fig. 1E).

### Identification of differentially methylated positions/regions (DMPs/DMRs) specific to CS

We took advantage of skin fibroblasts derived from very rare UVSS patients (that do not display the progeroid phenotype) to identify DNA methylation changes specifically related to precocious ageing in CS, and select out the impaired response to UV. For this, we compared genome-wide DNA methylation profiles between WT and UVSS cells, merged in the same “Non-Aged” group, *versus* CS derived-fibroblasts (“Aged” group). We first searched for differentially methylated positions (DMPs) between the Non-Aged and Aged samples (Fig. S1). Out of 441,982 analysed probes, we unveiled 15,648 DMPs (ANOVA; BH-corrected *p*-value <0.05; 3.5% of total probes), 13,177 of which were hypomethylated and 2,471 hypermethylated in the Aged group (top-ranking hits in Table 1; Fig. 1F; Table S1). About one third of DMPs (5,412) mapped in a genic sequence. Results of the Fisher’s exact test to assess whether DMPs are over- or under-represented in specific genomic regions revealed that both hypo- and hyper-methylated DMPs were under-represented (p-value<0.05) in Islands (DNA segments of >200bp in length, with high CpG density, >50% GC percentage, >60% observed/expected CpG ratio). Hypomethylated DMPs were strongly enriched in Open Sea (isolated CpGs in the genome) and Shelves, whereas hypermethylated DMPs were enriched in Shores, Shelves and Open Sea regions (Fig. 1G). Unsupervised hierarchical clustering on the list of 15,648 DMPs confirmed a clear separation between the Aged and Non-Aged groups (Fig. 1H). Furthermore, CS subtypes tended to cluster in separate subgroups, such as WT and UVSS cells.

**Table 1.**
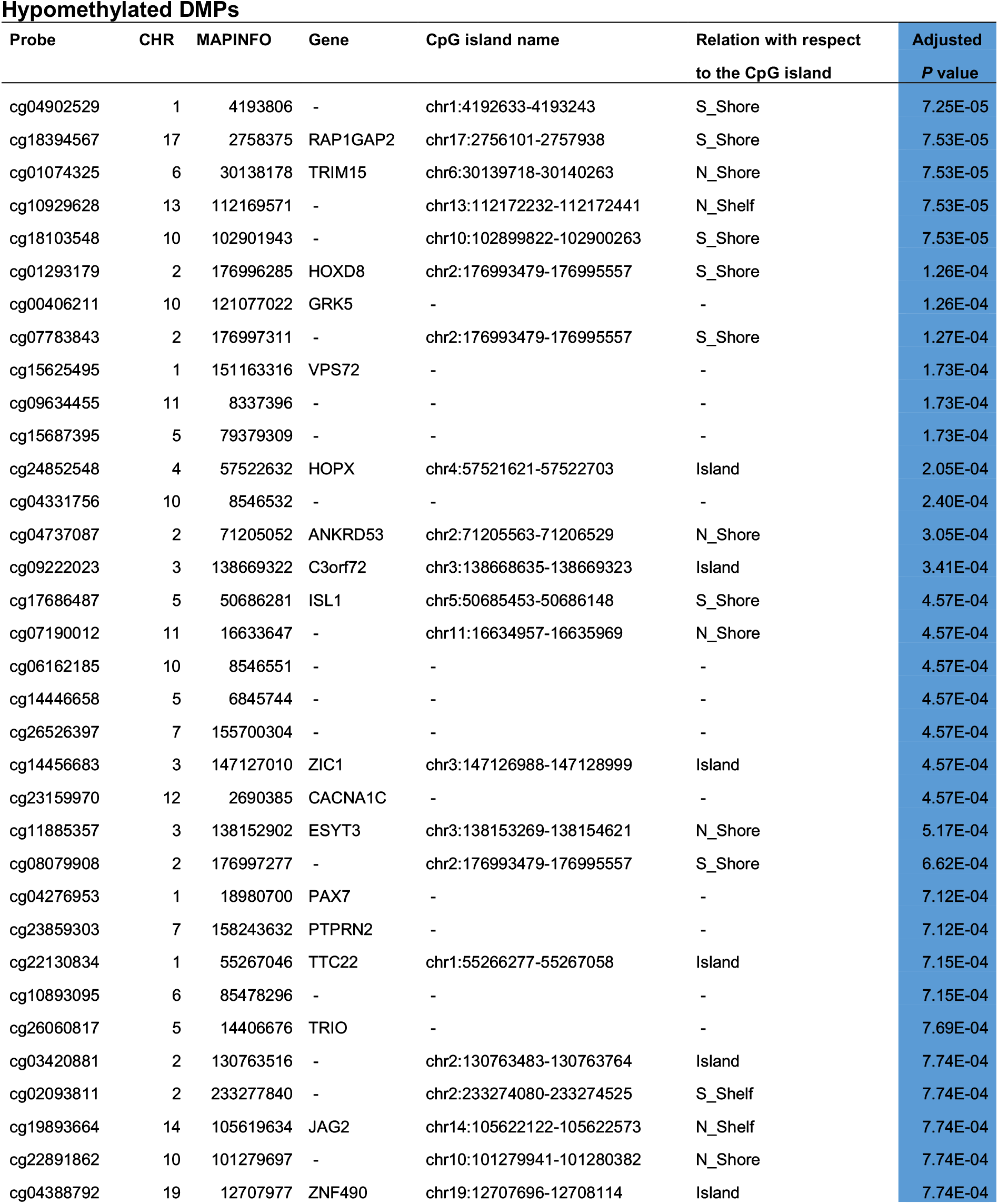
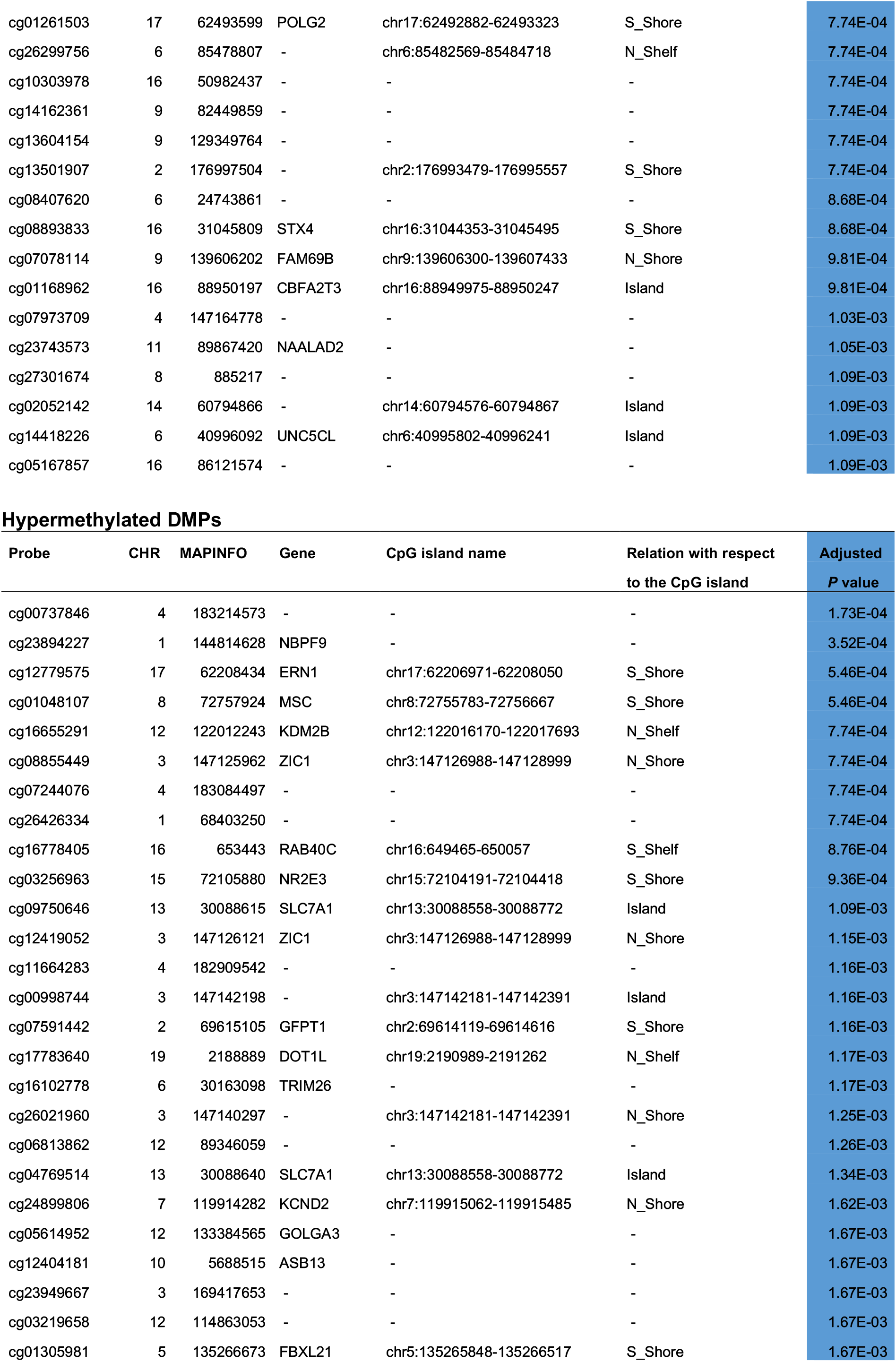
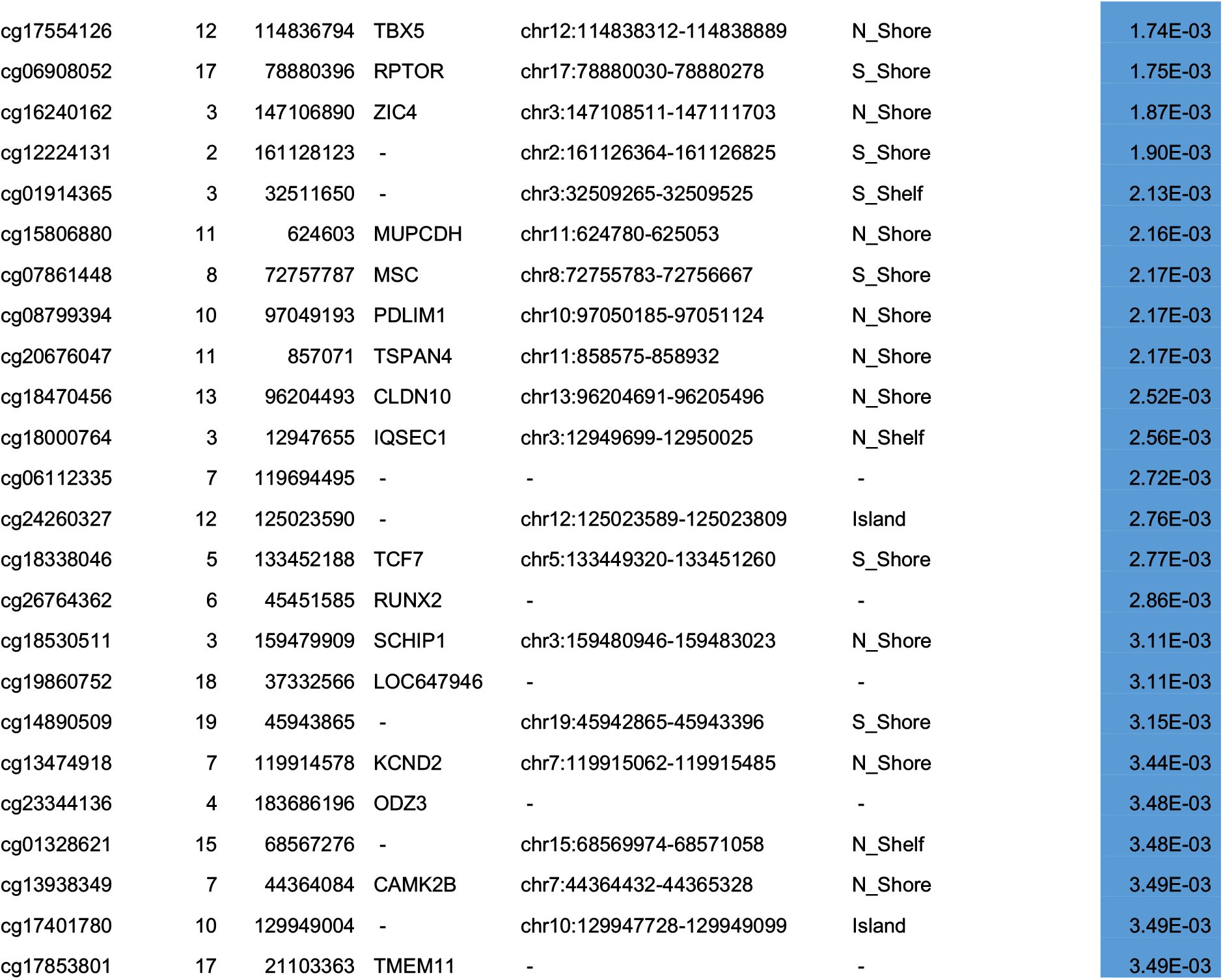
Top 50 hypo- and hyper-methylated DMPs of Aged *versus* Non-Aged condition upon single CpG analysis. DMPs are ranked according to the adjusted p-value. Mapping information on DMPs are given by the probe ID, the chromosome and base pair numbers, the CpG Island coordinate, the relation respect to the CpG island, and the associated gene (when applicable).

To strengthen the analysis of genes associated with CS-specific CpGs methylation changes, we searched for differentially methylated regions (DMRs). We focused on probes located on CpG Islands and surrounding regions (Shores and Shelves) of gene-associated regions and applied a pipeline based on a Multivariate Analysis of Variance (MANOVA) of adjacent probes, which has been validated for relevant age-associated changes^32^. We identified 1817 DMRs in Aged *versus* Non-Aged groups, of which 1416 were hypomethylated and 401 hyper-methylated (MANOVA BH-corrected *p*-value <0.05; Table S2, sheets 1 and 2). The top-ranking hits of DMRs list are reported in Table 2. These 1817 DMRs map in 1498 genes (1140 of which are hypomethylated and 358 hypermethylated), since some genes are covered by multiple DMRs.

**Table 2.**
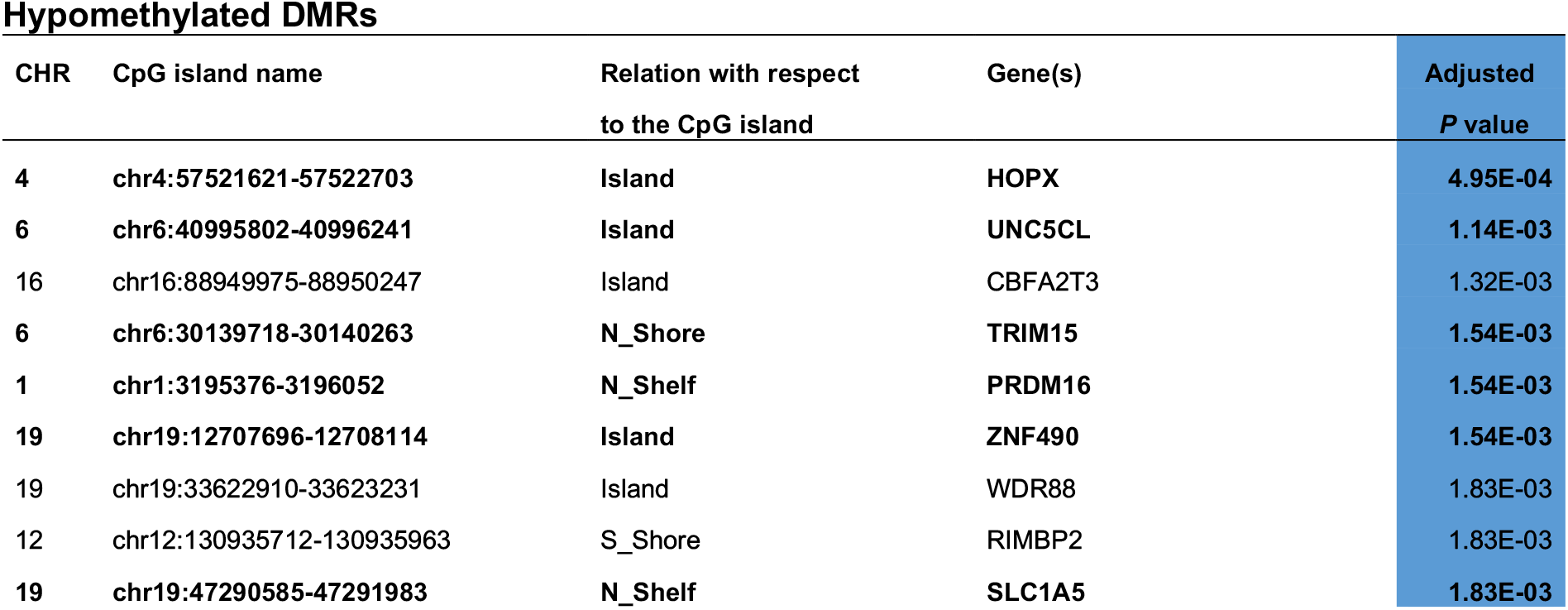
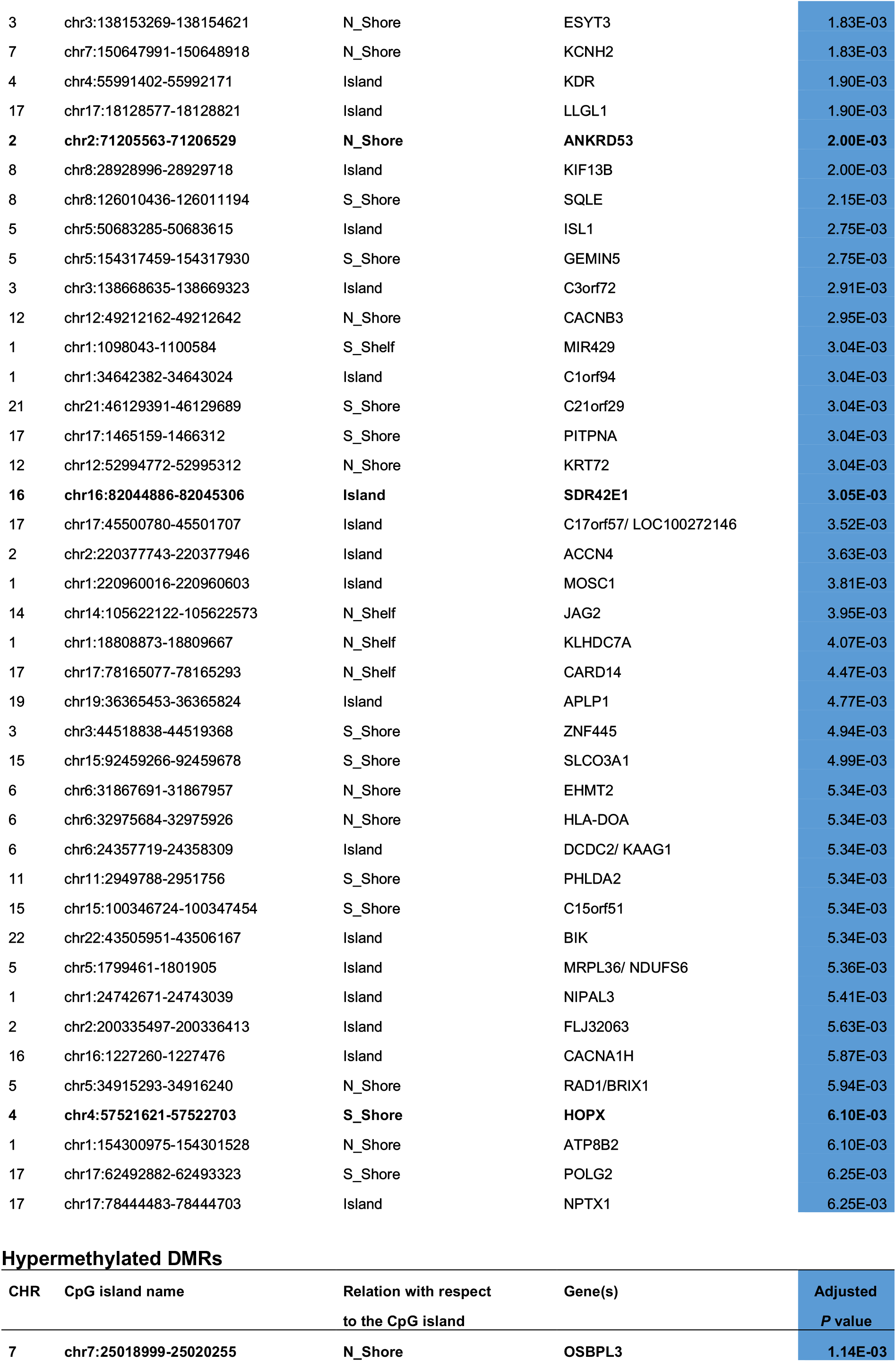
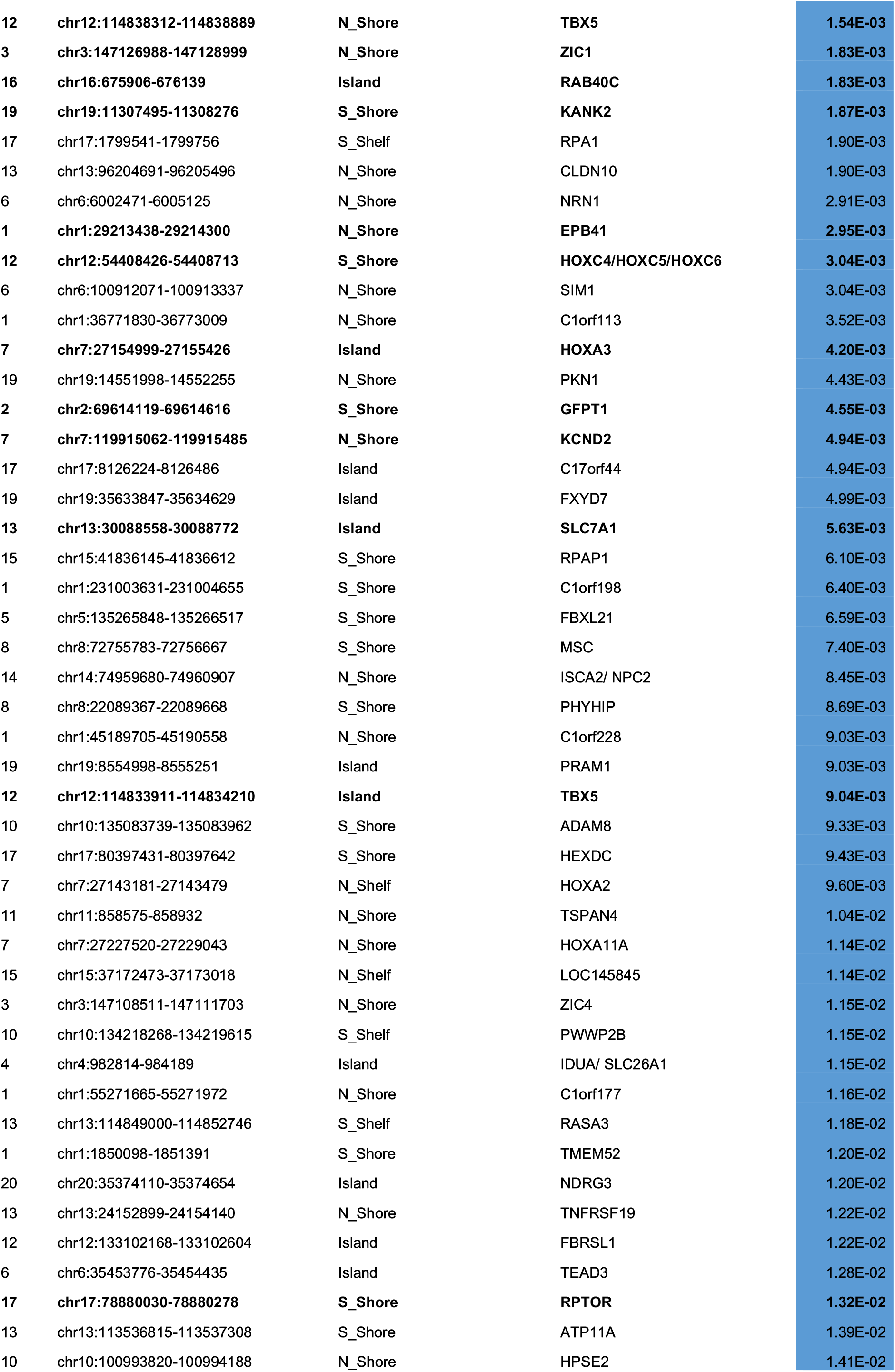
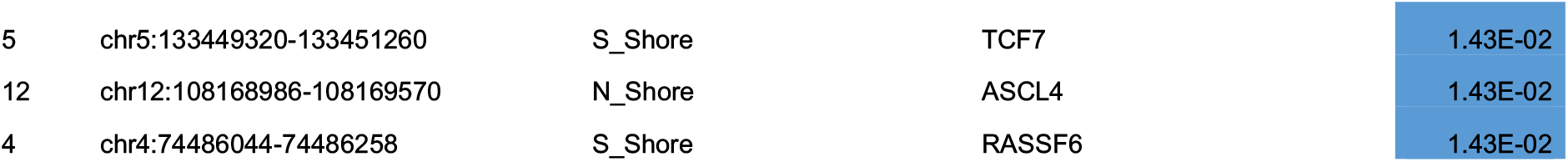
Top 50 hypo- and hyper-methylated DMRs of Aged *versus* Non-Aged. DMRs are ranked according to the adjusted p-value. Mapping information on DMRs are given by the chromosome number, the CpG Island coordinate, the relation respect to the CpG island, and the associated gene. Lines in bold indicate DMRs and corresponding genes assessed for expression.

Importantly, in most DMRs, UVSS samples displayed an intermediary DNA methylation profile between WT and CS samples (Fig. S3A) rather than being close to WT (Fig. S3B). We reasoned that some DNA methylation changes in this list could be linked to impaired DNA repair (a feature common to UVSS and CS cells) rather than precocious ageing (only in CS). We thus filtered the DMRs list in order to retain only those that unambiguously separated Aged (CS) samples and Non-Aged samples, thus having the highest probability of being implicated in the mechanism leading to the precocious ageing phenotype (see Materials and Methods). This approach resulted in the exclusion of 88% of the hypo- and 87% of the hyper-methylated previously identified DMRs and established a shorter and more stringent list of 222 hits, that we named stringentDMRs (Fig. S1 and Table S3), including 141 hypo- and 81 hyper-methylated regions in Aged compared to Non-Aged. The top-ranking hits of this list are reported in Table 3, together with a description of the function of the associated genes and of the pathological condition(s) in which they have been implicated. The implication in pathology was available for 33 out of the 40 most hypomethylated/hypermethylated genes, and revealed that 18/33 genes are involved in neuromuscular alterations and 9/33 in cancer. Hierarchical clustering on the stringentDMRs list confirmed a clear separation between the Aged and Non-Aged group, a clear clustering of CS subtypes, UVSS and WT cells in separate subgroups (Fig. S3C), and the maintenance of top-ranking position of individual genes compared to the non-filtered DMRs list. Altogether the 20 most hypomethylated and 20 most hypermethylated genes code for a microRNA, a long intergenic non-coding region, and 9 genes of unknown function, and the remaining 29 genes code for transcription factors (11/29), transporters (7/29), proteins involved in metabolic pathways (4/29), cytoskeleton dynamics (2/29), and epigenetics (2/29) (see Table 3).

**Table 3.**
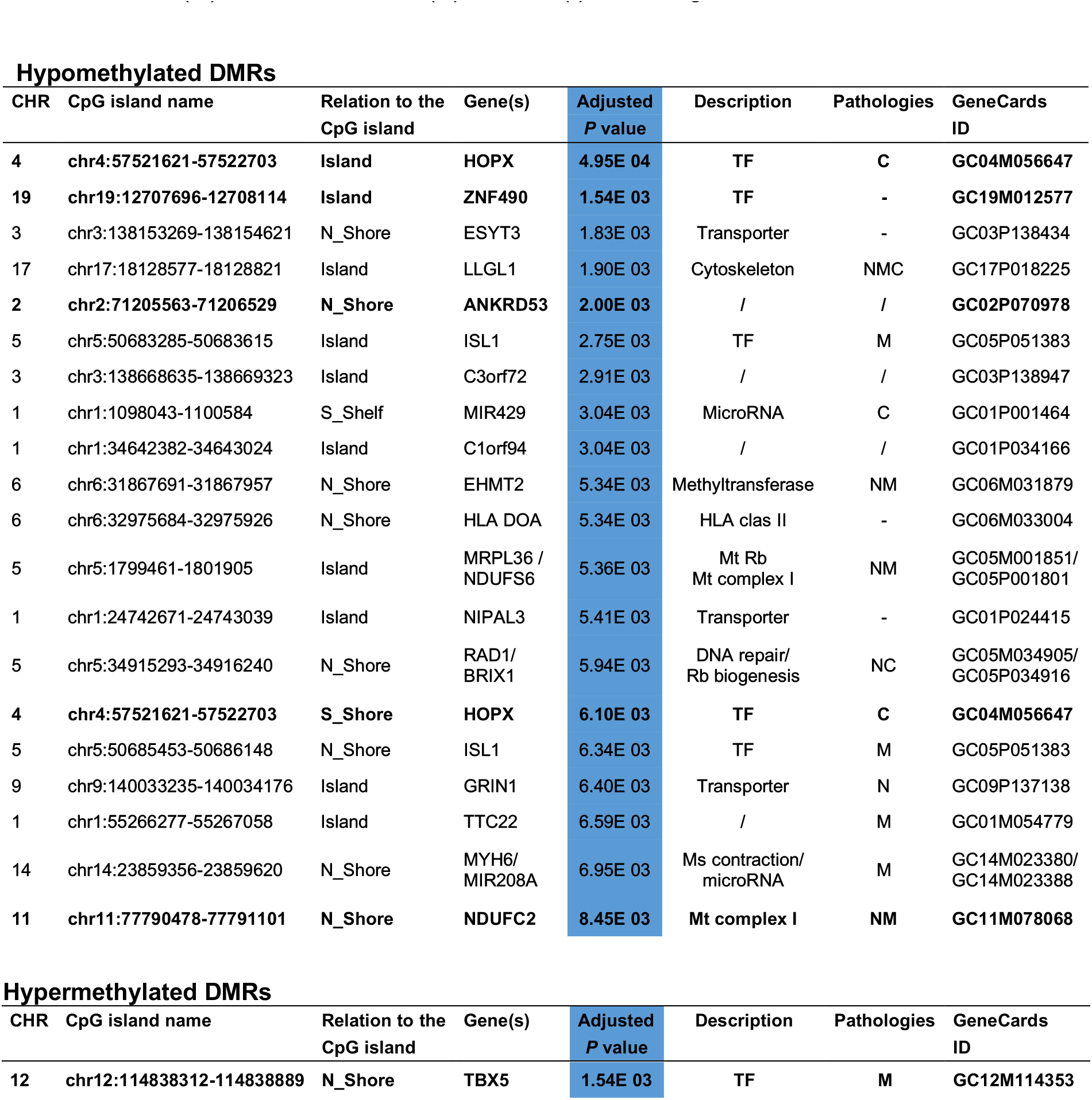
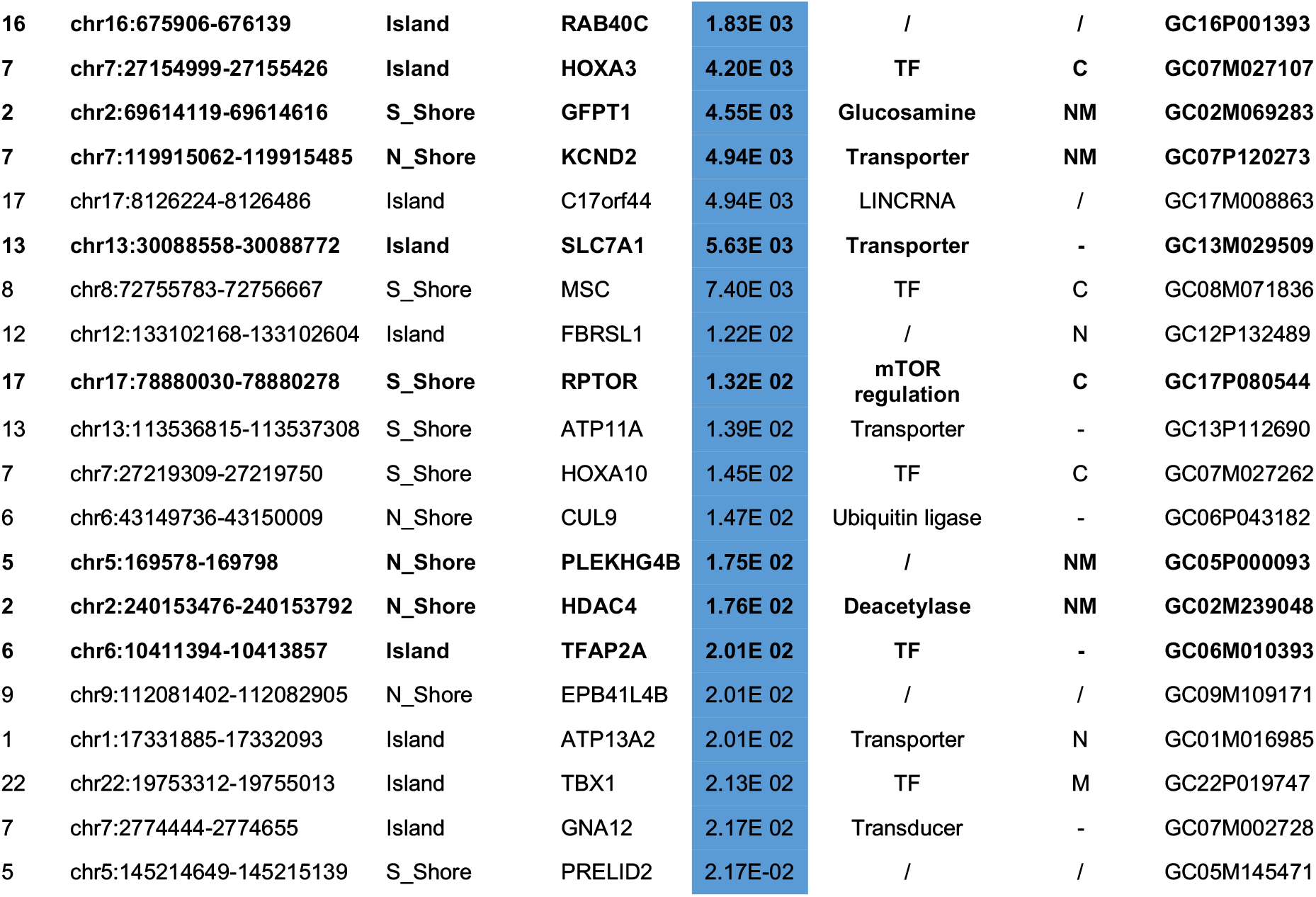
Top 20 hypo- and hyper-methylated stringentDMRs of Aged *versus* Non-Aged. DMRs are ranked according to the adjusted p-value. Mapping information on DMRs are given by the chromosome number, the CpG Island coordinate, the relation respect to the CpG island, and the associated gene. A description and the GeneCards ID of each gene are also provided. Bold lines indicate stringentDMRs and corresponding genes assessed for expression. TF: Transcription factor; Mt: mitochondrial; Rb: ribosome; Ms: Muscle. “/” is unknown. Implication in pathological states that affect the neurological (N) and/or muscular (M) function, or cancer (C), or none (-), according to GeneCards ID.

### Functional enrichment analysis of methylated positions and regions in CS

To characterize the potential biological impact of DNA methylation changes in CS, we performed a gene set enrichment analysis (GSEA) on the genes associated with methylated positions (MPs) and regions (MRs) resulted from the Aged *vs* Non-Aged comparison, using the Genome Ontology (GO) database. GSEA is a method that determines whether an a priori defined set of genes shows statistically significant concordant differences between two biological states (here Aged *versus* Non-Aged). Since both MPs (methylated positions) and MRs (methylated regions) results displayed high semantic redundancy of significantly enriched GO terms (*i.e*. Adjusted p-value<0.01, not shown), we removed the dispensable terms using REVIGO^33^. The treemap visualisation of the remaining terms, which allows hierarchical grouping of semantically related terms, showed that the largest identified “superclusters”, *i.e*. genes implicated in embryonic morphogenesis, transport of ions, and regulation of the membrane potential of synapse, were shared in between MPs (Fig. S4A) and MRs (Fig. S4B). This indicates that genes associated with MPs and MRs in CS are functionally comparable and underscores the robustness of our analyses.

We then focused on the functional analysis related to MRs, that we assume to be more prone to influence the phenotype than MPs. We identified 47 GO terms significantly associated with methylation changes between Aged and Non-Aged samples (Fig. 2A). These GO terms include 32 biological processes, 6 molecular functions, and 9 cellular components that in total contained 2100 genes (Table S4, sheet 1), 580 of which were associated with DMRs also identified in CS (Fig. 2B, listed in Table S4, sheet 2).

**Figure 2.**
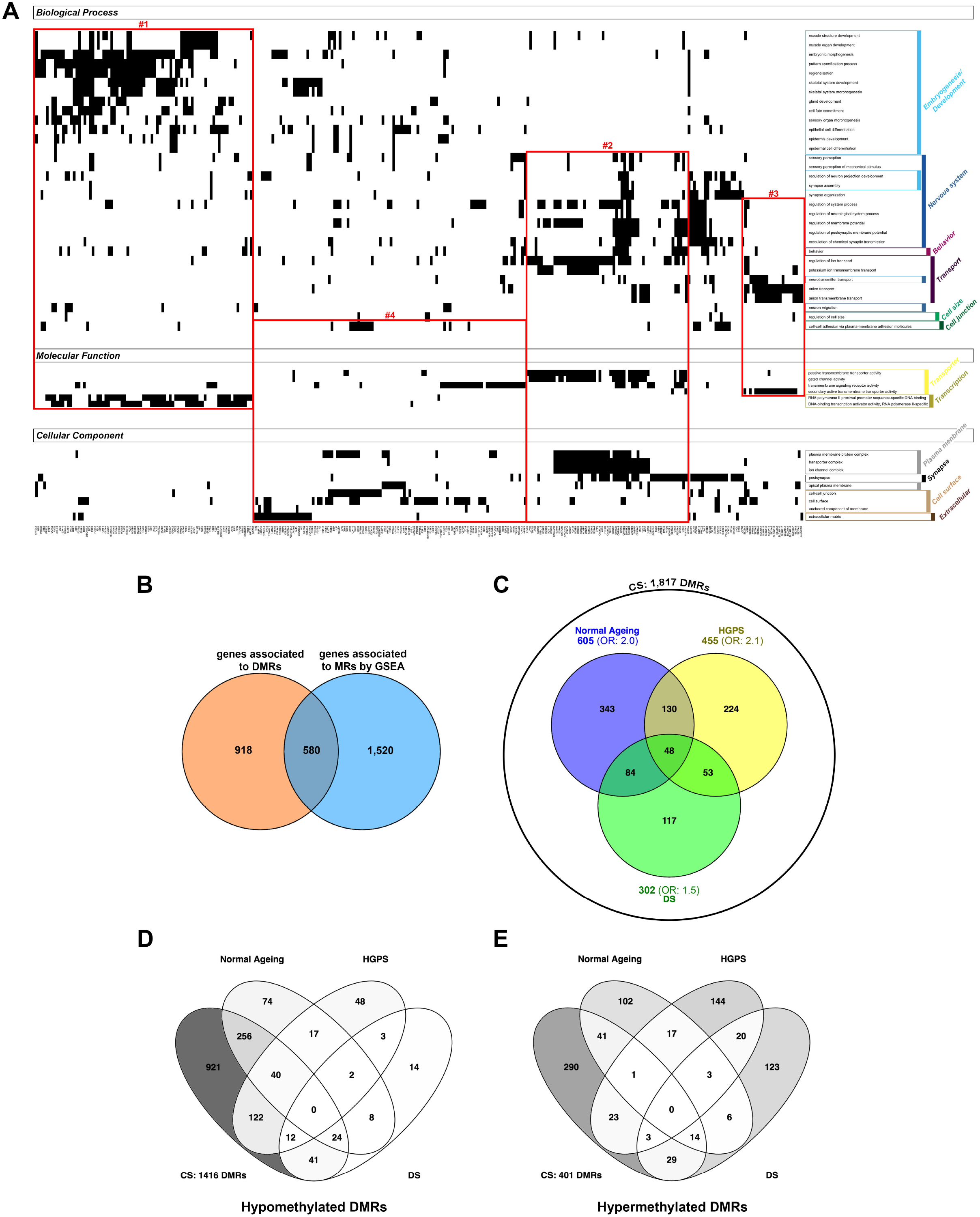
Filtering and functional GO enrichment analysis (GSEA) of methylated regions in Aged *versus* Non-Aged. (**A**) Plots reporting the correspondence between the enriched GO terms identified in the Aged *vs*. Non-Aged GSEA analysis (right y-axis) and the genes associated to DMRs, X-axis. This representation displays genes in the X-axis, and their corresponding GO term annotation on the Y-axis. GO terms on the Y-axis are grouped by GO categories (biological process, molecular function, cellular component). Genes may belong to more than one GO category as well as more than one GO term. Indeed, the same gene can be annotated with in multiple semantically similar terms, and the three GO categories provide complementary biological information. Numbered red frames indicate identified clusters of genes (related GO terms/GO categories); #1: transcription factors implicated in embryogenesis/development, #2: transmembrane transporters/channels implicated in the nervous system, #3: active ions/neurotransmitters transporters, #4: components of the extracellular matrix or cell surface possibly implicated in cell junction/adhesion or in signal transduction. (**B**) Proportion of genes responsible for the GO enrichment (580) in the total list of genes associated to DMRs (1498) and out of the total genes identifies by the GSEA analysis (2100). Venn diagram showing (**C**) the number of common and specific DMRs in Normal Ageing, HGPS and DS datasets, among the 1817 DMRs identified in CS (represented with an external diagram) and the common and specific hypo- (**D**) and hyper-methylated (**E**) DMRs in CS, Normal Ageing, HGPS, and DS datasets.

We summarised the functional relevance of the 580 DMR-associated genes by simultaneously clustering every single genes according to their GO annotation and GO terms by their semantic proximity (Fig. 2A). We observed that DMRs-associated genes are principally enriched in transcription factors implicated in embryogenesis/development (Red frame #1 in Fig. 2A), transmembrane transporters/channels in the nervous system (post-synapse regulation) (Red frame #2), active ions/neurotransmitters transporters (Red frame #3), and components of the extracellular matrix or cell surface, possibly implicated in cell junction/adhesion or in signal transduction (signalling receptors) (Red frame #4). Genes present in only one of the three categories (biological process or molecular function or cellular component) were regrouped separately (Fig. S4C). These last clusters included genes involved in muscle, skeletal, epidermis, and synapse development, as well as cell size regulation (Red frames #5, 6, 7, 8 and 9 in Fig. S4C, respectively), suggesting that they have a more tissue-specific function than those reported in Fig. 2A.

### Comparison of CS-specific DMRs with other progeroid syndromes and normal ageing

To assess whether CS-specific changes in DNA methylation are shared in normal ageing and other progeroid diseases or, conversely, CS changes display a specific epigenetic signature, we assessed available Infinium datasets analysing: 1) other progeroid syndromes: HGPS (fibroblasts from unpublished dataset, see Materials and Methods), WS (whole blood)^26^, and DS (whole blood)^25^, and 2) mesenchymal stem cells (MSC) from individuals of different age, as a proxy of age-associated changes in human fibroblasts^34^. We first compared the 15648 CS-DMPs (this study, Table S1) with DMPs identified in the other datasets. As summarized in Table 4, out of 15648 CS-specific DMPs, 2196 (14%) were also detected in the HGPS dataset, 1 in the WS dataset, 1004 (6%) in the DS dataset and 2286 (15%) in the MSC ageing dataset. With the exception of the WS-DMPs, CS-specific DMPs were significantly enriched in DMPs identified in the other experimental models (p-value <0.05, Fisher’s Exact Test; odds ratio of 1.6, 1.5 and 1.6 for HGPS, DS and MSC datasets, respectively). The direction of epigenetic changes (hyper- or hypomethylation compared to the respective controls) was consistent with that observed in CS for about 50% of DMPs in other progeroid diseases (except DS, 33.5 %), and raised to 69.70% for DMPs in common between CS and normal MSC aging (Table 4).

**Table 4.**
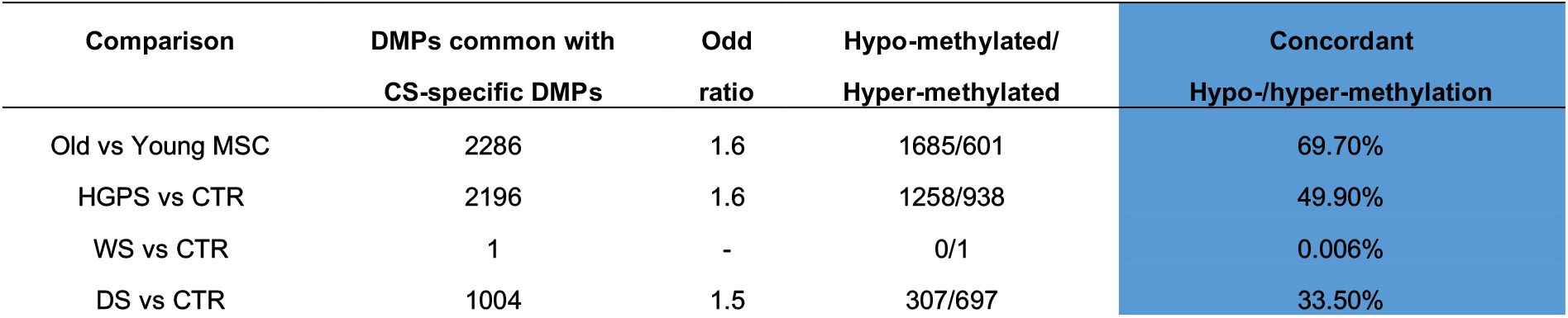
Comparison of the CS-specific DMPs with those emerged in other progeroid conditions (HGPS, WS and DS) and in normal aging of MSC. Number of common DMPs between DMPs identified in Aged vs Non-Aged (15648) conditions and DMPs identified in other studies performed in progeroid syndromes or during normal ageing. For each, the odd ratios of Fisher’s Exact Tests (p-value <0.05), the number (out of total) and percentage of concordant hyper-/hypo-DMPs (same direction) compared to the respective controls is also provided.

We then confronted the 1817 CS-specific DMRs with DMRs identified in the other datasets (Fig. 2C, Table S2, sheet 2). No DMRs were identified in WS using the same significance threshold applied to the other datasets. Conversely, 25% (455 DMRs), 17% (302 DMRs) and 33% (605 DMRs) of CS-specific DMRs were also significant in HGPS, DS, and normal ageing, respectively (Fig. 2C). These overlaps were greater than expected by chance (p-value <0.05, Fisher’s Exact Test; odds ratio of 2.1, 1.5 and 2 for HGPS, DS and MSC datasets respectively). Forty-eight DMRs were common to all datasets (not considering WS) and are highlighted in Table S2 (sheet 2). Differently from DMPs, the classification of DMRs as hyper- or hypo-methylated is not a straightforward task, as the same DMR can include both probes that gain and probes that loss methylation in cases compared to controls. We attempted to assign the DMRs as hypermethylated or hypomethylated by considering the methylation changes of the most significant CpG probe within each DMR. The results reported in Fig. 2D and E show that the normal MSC ageing has the highest proportion of DMRs concordant with CS, similarly to what observed for DMPs. None of these 48 DMRs were found to change in the same direction in all categories (CS, HGPS, DS, and normal ageing) (Fig. 2D and E).

### The methylation clock

We analysed our dataset with the “skin and blood clock” (based on 391 CpGs), which was explicitly designed for *in vitro* studies with fibroblasts^29^. By contrast, the original pan tissue clock from Horvath (2013) is less accurate for human fibroblasts^22^. The skin and blood clock showed that CS samples exhibit substantial epigenetic age acceleration: four CS-I children and three CS-II newborns displayed an average DNA methylation (DNAm) age of 28.8- and 7.7-year-old, respectively (Table 5). Conversely, WT and more importantly UVSS patients did not show signs of accelerate epigenetic ageing, in the sense that the age estimate, DNAm age, was close to the chronological age of the donors/patients.

**Table 5.**
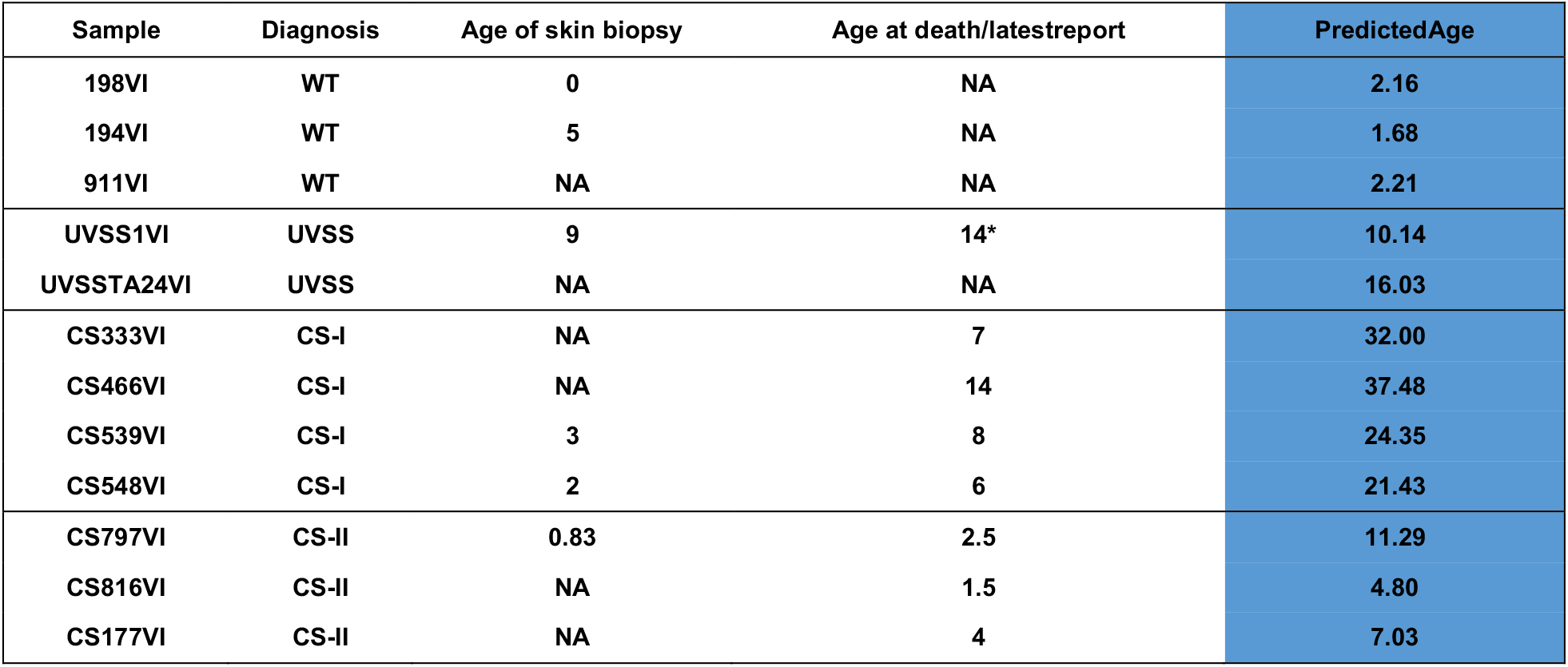
Methylation clock. For each cell type is shown the age at skin biopsy, age at death/age at latest follow up. The star *, indicates that the individual was alive at the last follow up (A. Sarasin, personal communication). DNAm Age refers to the predicted age based on the skin and blood clock (Horvath 2018). All age columns are in units of years. NA: Not available.

### Correlation between DMRs and gene expression

Transcriptomic data were not available from our samples, we therefore assessed by RT-qPCR the expression of several key genes identified as differentially methylated in CS. We selected DMRs according to one or more of the following criteria: the lowest *p*-values, the lowest deltas of methylation between WT and UVSS, and also their presence in related datasets (normal ageing and/or other progeroid diseases, see Fig. 3). Fourteen of the selected genes were included in the stringentDMRs list. The correlation between DNA methylation and transcription for each cell sample and for each of the 31 selected genes is shown in Figure 3. Fifteen genes showed a strong (0.4 ≤ R2 ≤ 0.9) and seven genes a moderate (0.4 ≤ R2 ≤ 0.2) correlation between DNA methylation and RNA expression. The majority of tested genes displayed an inverse methylation/transcription correlation whereas 12 DMRs were poorly or not correlated with changes in gene expression. Importantly, significantly relevant inverse correlation was observed for 8 genes (*EPB41*, *EPB49*, *ASAH1*, *HOXA11*, *VARS*, *SLC1A5*, *PRDM16*, and *ZIC1*), and direct correlation for 3 genes (*SCLA7A1*, *NDFUC2*, and *CLDND1*) that include genes common to multiple progeroid diseases and normal ageing.

**Figure 3.**
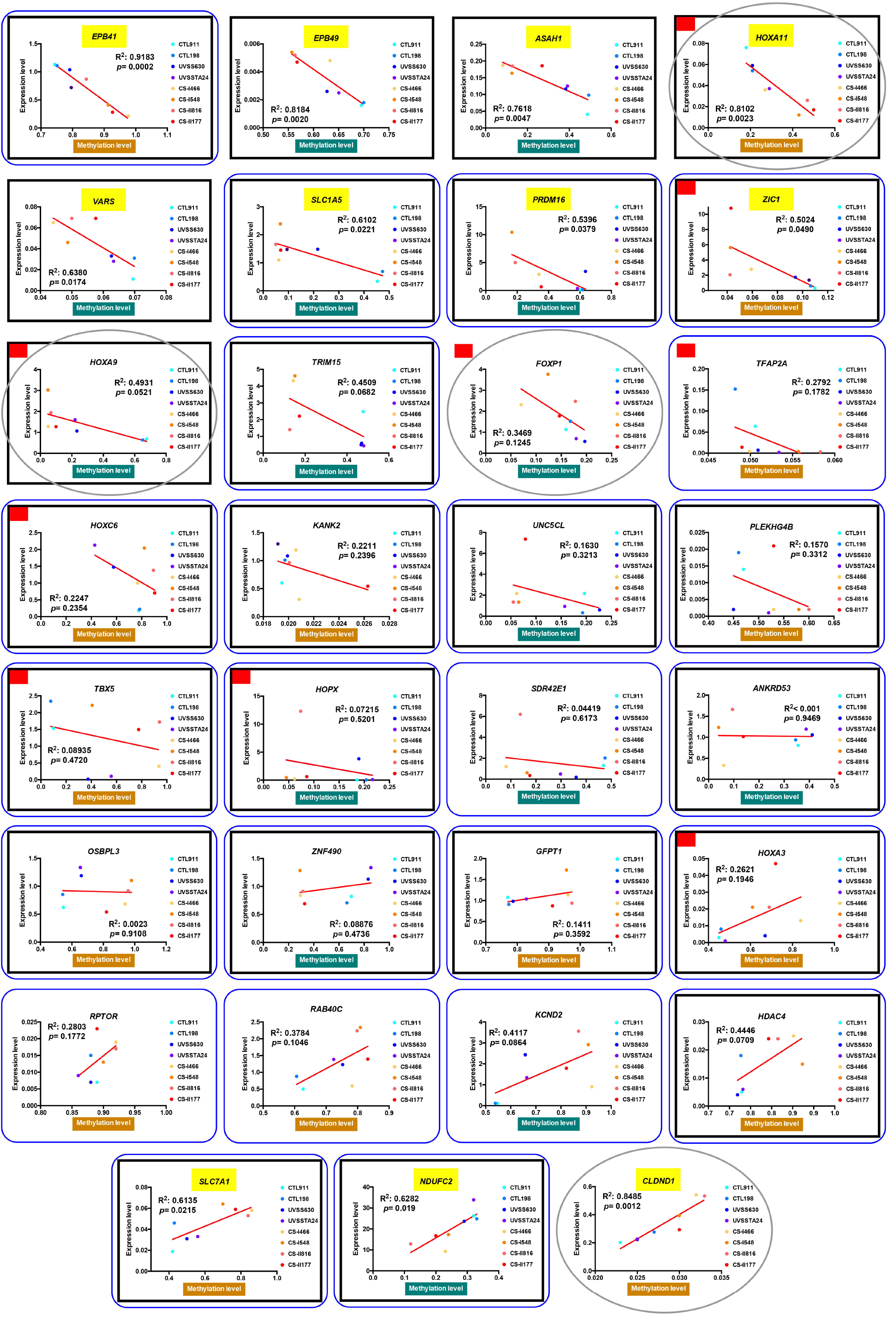
Correlation between DNA methylation and transcription levels in CS cells. For each cell type, the expression of selected genes was plotted against the methylation level of their DNA. The linear regression curve (Graphpad Prism) is indicated in red. Pearson’s Correlation Coefficient (Graphpad Prism) was used to assess the correlation between methylation and transcription levels. R squared (R^2^) and *p-value* (*p*) are indicated in each graph. The correlation was considered significant when p<0.05. Underscored gene name (yellow) indicates significant correlation. Nineteen out of 31 genes tended to be negatively correlated, 8 of which were significantly relevant (*EPB41*, *EPB49*, *ASAH1*, *HOXA11*, *VARS*, *SLC1A5*, *PRDM16*, and *ZIC1*). Four other genes were non-correlated (*ANKDR53*, *OSBPL3*, *ZNF490* and *GFPT1*), and 8 genes tended to be positively correlated (*HOXA3*, *RPTOR*, *RAB40C*, *KCDN2*, *HDAC4*, *SLC7A1*, *NDUFC2*, *and CLDND1*). The characteristics of selected genes are indicated below. Gold and green frames indicate the direction of methylation changes (hypermethylation and hypomethylation in the Aged vs Non-Aged group, respectively). Genes that showed differential methylation also in normal ageing (*EPB49*, *ASAH1*, *HOXA11*, *VARS*, *SLC1A5*, *PRDM16*, *HOXA9*, *HOXC6*, *KANK2*, *UNC5CL*, *PLEKHG4B*, *TBX5*, *HOPX*, *ANKRD53*, *OSBPL3*, *HOXA3*, *HDAC4*, *SLC7A1*, *NDUFC2*) and/or other progeroid diseases (*EPB41*, *EPB49*, *ASAH1*, *HOXA11*, *VARS*, *SLC1A5*, *PRDM16*, *ZIC1*, *HOXA9*, *TRIM15*, *HOXC6*, *KANK2*, *UNC5CL*, *PLEKHG4B*, *TBX5*, *HOPX*, *ANKRD53*, *OSBPL3*, *ZNF490*, *GFPT1*, *HOXA3*, *HDAC4*, *SLC7A1*, *NDUFC2*) are framed in black. Genes present in the top list of DMRs (*EPB41*, *SLC1A5*, *PRDM16*, *ZIC1*, *TRIM15*, *HOXC6*, *KANK2*, *UNC5CL*, *TBX5*, *HOPX*, *SDR42E1*, *ANKRD53*, *OSBPL3*, *ZNF490*, *GFPT1*, *HOXA3*, *RPTOR*, *RAB40C*, *SLC7A1*, *NDUFC2*) and/or stringentDMRs (*TFAP2A*, *PLEKHG4B*, *TBX5*, *HOPX*, *ANKRD53*, *ZNF490*, *GFPT1*, *HOXA3*, *RPTOR*, *RAB40C*, *KCND2*, *HDAC4*, *SLC7A1*, *NDUFC2*) are framed in dark blue. Genes selected specifically for their function (*HOXA11*, *HOXA9*, *FOXP1*, *CLDND1*) are framed in grey (ovals). Genes selected to be specifically differentially methylated in CS are *FOXP1*, *TFAP2A*, *SDR42E1*, *RPTOR*, *RAB40C*, *KCND2*, and *CLDND1*. These characteristics/criteria are not mutually exclusive. TFs are identified with a red square.

## Discussion

The cause of premature ageing features in Cockayne syndrome remains largely unknown despite the underlying mutations are known since decades. Increasing evidences suggest that the DNA repair defect alone does not explain the progeroid phenotype in CS^35^. In this context, previous work from our group revealed an increase in the expression of the HTRA3 protease as a trigger of CS-specific defects (which are absent in UVSS) leading to mitochondrial dysfunction^17^. Since HTRA3 expression is regulated by DNA methylation in some cancers^36^, we wondered whether CS is associated with genomewide DNA methylation changes. Importantly, our analysis relies on the methodological assumption that DNA methylation changes that are relevant for the progeroid phenotype of CS are not shared with UVSS. The results that we achieved support this assumption, suggesting that a specific methylation signature characterizes CS and contributes to its progeroid phenotype.

Unsupervised analysis of DNA methylation profiles in our dataset revealed that samples cluster together according to the classification of the disease. This is of particular relevance since CS patients display a large clinical heterogeneity, which constitute almost a continuum of severity of features^37^, and have obviously distinct genomic backgrounds. Notably UVSS samples, that carry the mutation but do not display features of CS, are closer to WT than CS samples. Moreover, within the same pathological class (CS type I), a clear separation was observed between cells mutated for *CSA* or *CSB* that are to date clinically indistinguishable^5^, and may therefore be useful to identify molecular differences in these patients. Curiously, CS type II cells displayed a larger spread in the PCA landscape, with some samples being close to UVSS, suggesting that qualitative differences play a major role in this condition.

CS cells tended to have lower DNA methylation values compared to UVSS or WT cells, resembling the global hypomethylation described in physiological ageing contexts and ascribed to heterochromatin loss in non-genic repetitive sequences with ageing^21^. By inferring the methylation of locus-specific repetitive elements from microarray data, we found that CS cells displayed a marked hypomethylation of Alu elements, while LINE elements were not affected by the syndrome. Thus, the methylation patterns of CS cells exacerbate those characteristic of normal ageing^38^. This result seems specific for our dataset, as in whole blood from WS patients no differences in DNA methylation of repetitive elements (experimentally measured) was found compared to healthy controls^39^, or this is due to differences between fibroblasts and blood cells. Interestingly, considering all the microarray probes and the inferred repetitive elements, DNA methylation of CS type I cells (mild form of the disease) was quantitatively more affected than CS type II cells (severe form of CS), suggesting that in addition to ageing-related dysfunctions other aspects of the pathology are associated with changes in the methylation state. Alternatively, DNA methylation changes at specific positions may have a more relevant role than others in the progeroid phenotype, thereby overcoming quantitative differences among samples (see below). We cannot rule out the possibility that methylation at some loci is a rescue mechanism hindering catastrophic effects, yet contributing to the ageing process.

To identify DNA methylation changes that could drive the precocious ageing phenotype of CS patients, we compared “Aged” (CS) *vs*. “Non-Aged” (WT+UVSS) samples in a differential analysis. We identified 15,648 DMPs in CS, 84% of which were hypomethylated, consistent with a study performed during regular ageing in cultured cells^34^ but differently from DMPs identified in progeroid WS and in the ageing blood, where only 45% and 46% of DMP were hypomethylated, respectively^26,40^. The consistency of CS-specific DNA methylation changes with regular ageing was further underscored by the observation that hypomethylated DMPs were enriched in isolated CpG (Open Sea) whereas hypermethylated DMPs were enriched in regulatory regions flanking gene promoters (Shore, Shelf)^41,42^. Thus, despite they represent the minority of DNA methylation changes hypermethylated DMPs are more prone to impact on gene transcription^21^ in CS.

To better evaluate the impact of DNA methylation changes on genes, we focused on DMRs encompassing genic sequences. This analysis identified 1817 DMRs in CS, 78% of which were hypomethylated. These DMRs cover 1498 genes, since some DMRs are present more than once in the same gene. We finally applied a filtering to retain only the DMRs where CS samples clustered separately from WT+UVSS, and that are reasonably specific to the progeroid/neurodegenerative phenotype (uncoupled from the DNA repair deficiency); overall view of the procedure in Figure S1. Using this pipeline and thanks to the pivotal role of UVSS samples (mutated but non-progeroid), we identified a CS-specific epigenetic signature highlighting a restricted list of 222 DMRs (64% hypomethylated). Successive steps of selection (from DMPs to DMRs and then to stringentDMRs) obviously restricted the number of hits but, importantly, the main gene categories remained highly represented along the successive analyses.

Indeed, the top 20 hypo- and hyper-methylated associated genes in the 222 stringentDMRs that constitute the CS epigenetic signature, in large part encode transcription factors implicated in developmental processes, membrane transporters (ions, amines, lipids), and cytoskeletal or membrane proteins involved in cell-to-cell contact. This list also included two subunits of the mitochondrial respiratory complex I (*NDUFS6* and *NDUFC2*). The expression of the membrane transporter *SLC7A1* (and to some extent also *KCND2*) and of the mitochondrial factor *NDUFC2* were directly correlated with DNA methylation levels in our samples (low expression in CS), suggesting functional relevance. A few hits in the stringentDMRs top list, *e.g*. the histone methyl transferase *EHMT2* (hypomethylated in CS) and the histone deacetylase *HDAC4* (hypermethylated in CS) have been previously linked to ageing^43,44^.

Analysis of pathways rather than single genes, using a gene set enrichment analysis (GSEA) and the gene ontology (GO) annotation, revealed GO terms that show statistically significant concordant differences between Aged and Non-Aged samples. Genes associated with DMRs were particularly enriched (high NES score) in the following biological processes/functions: i) developmental transcription factors and regulators, ii) ion transporters, and iii) cell junction factors, in remarkable agreement with single gene analyses above.

i. Gene enrichment in embryogenesis/developmental transcription factors (TFs) prevails over all other GO terms, suggesting a major implication in CS-specific defects, and to a larger extent than observed in other precocious ageing diseases^25^ and during regular ageing^40^. Several developmental TFs, which are differentially methylated in CS, have been linked to ageing-related processes, as the homeobox gene *HOXA9*^45^, TFs of the T-box (*TBX1*, *4*, *5*, *15*) and Fox (*FOXA1*, *FOXF1*) families, Wnt transcription regulators (*WNT7A*, *WNT8B*, *FDZ1*, *FDZ2*, *FDZ7*), and GATA, SOX, and PAX family members^46–48^. These examples underscore the relevance of the CS model for ageing. To note, a fraction of these genes, including HAND (*HAND1*, *2*), LHX (*LHX2*, *3*), NKX (*NKX2-5*, *2-8*, *6-1*), and PITX (*PITX1*, *2*) families, to our knowledge have not been associated with ageing, possibly being novel developmental regulators factors that play a role in CS as well in regular ageing. *HOXC6*, *HOXP*, and *HOXA3* DNAm levels did not show a significant correlation with gene expression, at least in the limited number of patient samples available. Other developmental transcription factors (*HOXA11*, *ZIC1*, and to some extent *HOXA9*) demonstrated a strong correlation between DNA methylation and expression levels, suggesting a functional impact of CS-specific methylations changes. These findings are consistent with the quasi-programmed theory of ageing that claims detrimental over-activity of developmental factors during ageing^49^. According to this theory if processes important early in life are not properly switched off in the adult, they progressively lead to aged-related alterations. Accordingly, developmental genes are normally tightly controlled and fail to be so during ageing^50^. We report modifications of DNA methylation in CS that are possibly responsible for deregulation of developmental genes, suggesting that this mechanism, whether or not of stochastic origin, plays a role in the ageing phenotype. Importantly, these TFs target multiple downstream genes and can thus directly drive the establishment of an aged-related transcriptional program. The identification of differentially methylated regions associated with genes implicated in tissue-specific development in CS (*i.e*. muscle or skeletal development) may provide clues to better understand why some organs are more susceptible to ageing than others^22,51^.
ii. Functional analysis of CS methylated regions reveals enrichment also in transmembrane ion transporters, both passive and active carriers. Passive transporters were mainly voltage-gated potassium channels (*KCNA3*, *KCNA4*, *KCNC2*, *KCND2*, *KCNE4*, *KCNG4*, *KCNH1*, *KCNH2*, *KCNIP4*, *KCNQ1*), which are implicated in regulating the membrane potential in excitable cells like neuronal or muscle cells^52^, but are also expressed in fibroblasts^53^. The involvement of these transporters in neurodegeneration has been evoked in several neurodegenerative diseases (*i.e*. Alzheimer’s and Parkinson’s disease)^54^, and they may play a role in neurodegeneration in CS. Interestingly, remodelling of K^+^ channels in dermal fibroblasts has been associated with the successful ageing of centenarians^52^, while overexpression of K^+^ channels has been observed in HGPS cells^55^. Several genes coding for ion channels identified as differentially methylated in our study such as GABA (*GABRD, GABRB3*), glutamate (*GRIN1, 2D*) or acetylcholine receptors (*CHRNA2, B2*), are of particular interest since they are also main neurotransmitter receptors that regulate brain function. Genes for active transmembrane transporters mainly involve the solute carrier group of transport proteins (SLCs) that carry a wide variety of substrates including ions, nutrients, and/or neurotransmitters through plasma/organelle membrane, including in fibroblasts^56^. Only a few members of this large family of transporters (> 400 proteins) have been associated with ageing^57^. *SLC1A5* and *SLC7A1* showed a strong direct or inverse correlation, respectively, with DNA methylation levels in CS patient and control samples.
iii. Gene enrichment in the functional analysis of CS methylated regions also revealed components of the cell surface and extracellular compartment, and included different types of cell adhesion, namely adherens junctions (*ACTN1, APC, CDH3, 4, 7, 15, JUP, DSP*), tight junctions (*CLDN6, 10*), and gap junctions (*GJA4, GJA8, GJC1*). The expression of *CLDN1* strongly correlated with DNA methylation levels in our samples. Cell adhesion factors play an important role in maintaining tissue architecture as well as tissue homeostasis due to their involvement in the formation of epithelial barriers that regulate exchanges, and is therefore not surprising that they are possibly involved in aging^58^. Accordingly, higher expression of protein involved in intercellular (tight) junctions was recently found in healthy centenarians compared to controls^59^. The role of these junctions, particularly during aging, remains poorly understood outside epithelia to which they seem restricted, despite fibroblasts express adhesion molecules^60^.

The CS epigenetic signature was compared with physiological ageing^34^ and other progeroid diseases (DS^25^, WS^26^, HGPS (unpublished data from GL)). Each progeroid disease is characterized by distinct features. For instance HGPS patients do not display neurodegeneration whereas CS patients do, and include neuronal loss, demyelination, gliosis and calcification, that unevenly affect the different parts of the CS brain^61^. As another example, WS patients display an increased propensity to develop cancer, whereas HGPS and CS patients typically do not develop cancer. Other clinical symptoms as growth impairment and skin abnormalities are shared among progeroid diseases and may result from common defective pathways.

Collectively, data from previous studies^25,27,28,39^ and our new analyses demonstrate that epigenetic remodelling is a common hallmark of progeroid syndromes. At the same time, as pointed out in the seminal study of George Martin^62^, commonalities and differences exist for changes occurring in progeroid disorders *versus* physiological aging. Our study contributes disentangling and clarifying the epigenetic basis of this insight. More specifically, we observed a significant overlap between DMPs and DMRs associated with CS and those identified in normal ageing, HGPS, and DS datasets. In particular, CS had 15% of DMPs in common with normal aging, which showed also the highest number of consistent DNA methylations changes (same direction) compared to other progeroid diseases. These results differ from those recently reported in WS whole blood cells, were only 6% of DMPs displayed age-associated DNA methylation changes^39^. Similarly, it was recently reported that epigenetic changes in HGPS fibroblasts differ from those occurring during physiological aging^28^. Our data thus suggest that CS DNA methylation changes are closer to normal ageing than other progeroid diseases, and confirm CS as a precocious ageing disorder. Importantly, the DNA methylation datasets that we analysed differ in size and derive from different tissues (fibroblasts for CS and HGPS, MSCs for normal aging, whole blood for DS and WS), a factor that may profoundly affect our results. Future studies should systematically analyse DNA methylation changes associated to progeroid disorders and to aging in comparable biospecimens.

Remarkably, several CS characteristics were distinct from normal ageing. For instance, significant hypomethylated regions were more numerous in CS than normal ageing, which rather displayed hypermethylated regions^40^. Moreover, in normal ageing hypermethylation was associated with developmental processes and DNA binding/transcription, whereas hypomethylation did not enrich a specific set of genes. Genes involved in transcriptional regulation of developmental processes, as indicated above, were prevalently hypomethylated in CS.

From the CS-specific pool, we identified 48 DMR-related genes that are common to the other data sets analysed (Normal Ageing, HGPS, DS). Among them, the expression of *EPB49*, *KANK2*, and to some extent also *HOXA9*, was inversely correlated with DNA methylation, suggesting that these methylation changes are relevant for CS. Several of these common genes are developmental TFs suggesting that epigenetically altered TFs are a common mark of other progeroid diseases and ageing, and not only CS. *EPB49* has been recently associated, together with other components of the haem metabolism, with human ageing^63^. Other genes in this list, like *NAGS*, a mitochondrial N-acetyl glutamate synthase linked to glutamine/glutamate metabolism, and GNL1, a little-known GTPase reported to modulate cell proliferation^64^, have not been associated with ageing to date, and the function of FAM43A is still unknown.

We applied to our dataset a methylation age predictor specifically developed for human fibroblasts^29^, which outperforms the pan-tissue age estimator^22^ and other clocks for this cell type. We showed acceleration of epigenetic age specifically in CS (but not in UVSS, as expected). Importantly, the methylation clock of CS was on average 3.5-fold larger for CS-I and up to 13.2-fold larger for CS-II, and these values are dramatically higher than in WS or DS (addition of 6.4 and 6.6 years, respectively, for 10-60 year old patients)^25,30^, suggesting that CS patient displays an extremely accelerated ageing. These data are in agreement with the accelerated aging we reported in six patients with CS (aged 3, 7, 8, 10 (twice) and 20 years) using the GlycoAgeTest, that measures the age-associated shift in serum N-glycan profile^65^. The fact that two distinct biomarkers of age (the epigenetic clock and the GlycoAgeTest) concordantly showed accelerated ageing in CS patients is remarkable, and mirrors what previously observed in DS, although at a much larger extent^25,66^.

A general trend in aged-related DNA methylation studies is that changes (hypo or hypermethylation) are not necessary correlated with changes in gene expression, even in regions with a highly reproducible epigenetic remodelling during ageing that define the “methylation clock”^22^. In the present study, we focused on DNAm changes located in gene regulatory regions. As available transcriptome data in CS involve conditions not compatible with our study, namely immortalized cells irradiated or not with UV^67^, or iPSC-derived neurons^68^, we rather analysed transcription levels of key genes directly in patient-derived fibroblasts. Transcription of 31 genes that displayed the highest DNA methylation changes in CS (top 50 hypo/hypermethylated genes) and/or were in common with other ageing datasets, unveiled a tendency to inverse correlation (high expression *vs* low methylation levels), although direct correlation was observed too, and one third of cases were statistically robust. The remaining cases showed no correlation or a tendency to direct correlation. The DNAm/transcription correlation appears remarkably larger than in other cases of ageing-related analyses (for instance less than 4% in normal ageing)^40^, perhaps as a result of the robustness of our data analysis (DMR instead of DMP, multiple filtering processes based on biological assumptions), or a characteristic of methylation changes in CS.

## Conclusions

We identified CS-specific DNA methylation changes by filtering out changes not associated with the progeroid and neurodegenerative phenotype, a unique characteristic of this pathological paradigm. Functional analyses revealed enrichment in developmental transcription factors, ion transporters, and cell junction factors, that appear thus critical for accelerated ageing/degeneration. In addition to epigenetic signatures in common between CS and physiological ageing, DNA methylations changes observed in CS may help identifying transcriptional changes that would appear at a later time in regular ageing. Finally, our results demonstrate that CS is a pertinent model to study ageing and unveil pathways and genes that could be relevant for future mechanistic studies.

## Material and methods

### Patient cells

Patient fibroblasts were derived from skin biopsies excised from unexposed body sites by dermatologists. Patients (or parents or legal guardians) provided informed consent to receive diagnosis and genetic analysis. The French Agency of Biomedicine (Paris, France) (Arrêté n°2001/904 and Ref: AG08-0321 GEN of 27/09/2008; (http://www.agence-biomedicine.fr/fr/experts/doc/praticiensGEN30092008.pdf) and the European Commission “Geneskin: Genetics of human genodermatosis” (Brussels, Belgium, 2008) approved this study. UVs24TA were a kind gift of dr. Graciela Spivak, Stanford University, USA. HGPS fibroblasts were from the BioLaM biobank at IGM-CNR, Bologna, Italy (IRCCS Istituto Ortopedico Rizzoli Ethical Committee approval no.0018250 – 2016).

### Cell culture conditions

Healthy, UVSS and CS cells (detailed in Fig. 1A) initially thawed at passage number (PN) 12 were cultured in Dulbecco’s modified eagle medium (DMEM; Gibco) supplemented with 2mM L-glutamine (GlutaMAX^™^; Gibco), 10% foetal bovine serum (FBS; Gibco), 1% Penicillin-streptomycin (Gibco) in 20% O2/5% CO2 at 37 °C. Cells were harvested at PN14 for further experiments.

### DNA extraction, bisulfite treatment and genome-wide DNA methylation analysis

Total genomic DNA (gDNA) from WT, UVSS and CS primary skin fibroblasts was isolated using the QIamp^®^ DNA mini kit (Cat#51304; Qiagen) and used for microarray-based analysis of genome wide DNA methylation levels in two different sets (see Fig.1A for detail of cells used for each set). For the first set, extracted gDNA was quantified by NanoDrop (ND-1000; Thermo Fisher Scientific) and 1μg was bisulfite-treated using the EZ DNA Methylation^™^ kit (Cat#D5001; Zymo Research), according to manufacturer instructions. Genone-wide DNA methylation analysis was performed on this material by Aros Applied Biotechnology A/S (Eurofins Genomics; Denmark) using the Infinium HumanMethylation450 BeadChip (Illumina). For the second set, extracted gDNA was directly send to Aros Applied Biotechnology A/S for bisulfite conversion and genome-wide DNA methylation using the Infinium MethylationEPIC BeadChip (Illumina) following manufacturer’s instructions.

The *minfi* Bioconductor package was used to extract raw signal intensities from each experimental set (Infinium HumanMethylation450 and Infinium MethylationEPIC BeadChips)^2^. All the samples were retained after preliminary quality checks (less than 1% of probes with a detection pvalue >0.05). The preprocessNoob function implemented in *minfi* was applied to normalize raw data. Common probes between the two experimental sets were then retained and annotated according to the Infinium HumanMethylation450 Beadchip annotation file. Methylation values were expressed as beta values (percentage of methylation, ranging from 0 to 1). Batch effects between the two experimental sets were corrected using the ComBat function implemented in the *sva* R package^69^ and, for the 7 samples replicated in the two experimental sets, mean beta values between the 450k and the EPIC experiment were calculated for each CpG probe. To predict the methylation of locus-specific repetitive elements (RE) we used the REMP bioconductor package^70^.

To identify DNA methylation differences between Aged and Non-Aged groups, probes on X and Y chromosomes were removed and the resulting dataset was analysed using: (1) analysis of variance (ANOVA), to identify differentially methylated positions (DMPs); (2) multivariate analysis of variance (MANOVA) on sliding windows of three chromosomally adjacent probes, as described in Bacalini *et al*.^32^, to identify differentially methylated regions (DMRs). The second approach was applied only to the probes mapping in CpG-rich regions (CpG islands, shores and shelves) associated to a gene, according to the Infinium HumanMethylation450 Beadchip annotation. Multiple testing correction was performed using Benjamini–Hochberg procedure. To identify the DMRs that unambiguously distinguished the Aged and Non-Aged groups, we further filtered the list of significant (BH-corrected pvalue <0.05) DMRs as follows: 1) for each DMR, we considered the sliding window of 3 adjacent probes having the smallest MANOVA p-value; 2) we performed k-means clustering on methylation values of the 3 probes, setting k=2 (corresponding to Aged and Non-Aged groups); 3) we retained only those DMRs in which all the CS samples were classified as Aged and all the WT and UVSS samples were classified as Non-Aged. Enrichment analysis was performed using Fisher exact test implemented in the R *stats* package. Heatmap and hierarchical clustering were performed using the *heatmap.2* function in R *gplots* package, with default settings (euclidean distance function, complete agglomeration method).

### Functional analysis

The functional scoring of the genes associated with MPs and MRs was done using Gene Set Enrichment Analysis (GSEA)^71^. The two biological samples were on one side the 7 Aged (CSI_A, CSI_B, and CSII) samples, and on the other side the 5 Non-Aged (WT + UVSS) samples. The specific implementation of GSEA for methylation datasets proposed by MethylGSA^72^ was used. MethylGSA adapts the robust rank aggregation (RRA) approach to adjust for number of CpGs in DNA methylation gene set testing.

The input for the GSEA analysis is the list of adjusted p-values per probe issued from the differential methylation analysis at the MR and MP level (see above). For the MRs, a representative probe is considered per region, the one holding the minimum adjusted p-value, leading to an initial set of 29603 probes. For MPs, instead, the initial set holds 220615 probes. Gene ontology terms (GO) with a minimum of 100 genes and a maximum of 500 genes associated were used to functionally score MRs. For MPs, KEGG pathways with a size varying between 5 and 500 genes were preferred.

For the visualization of GO terms associated with significant methylation changes in MRs and their corresponding genes (Fig. 2A), a selection of terms and genes was done. The list of significant GO terms (GSEA adjusted p-value<0.1) was simplified by removing redundant terms with the REVIGO procedure^33^ with default parameters and keeping only genes overlapping with genes associated to DMRs.

### Comparison with other methylation datasets

Publicly available Infinium450k (GSE52114 and GSE52588) and InfiniumEPIC (GSE100825) datasets were downloaded from the GEO database. GSE52114 includes DNA methylation profiling of mesenchymal stem cells obtained from 22 young individuals (age range 2-29 years) and 12 old individuals (age range 61-92 years). GSE52588 includes DNA methylation profiling of whole blood DNA from rom 29 subjects affected by Down syndrome (age range 10-43 years), their unaffected siblings (age range 9-52 years) and their mothers (age range 41-83 years). GSE100825 includes DNA methylation profiling of 3 subjects affected by Werner syndrome (age range 44-53 years) and three age- and sex-matched healthy controls. Finally, we used an unpublished Infinium450k dataset which includes DNA methylation profiling of fibroblasts from one HGPS patient (6 years; 3 replicates assessed) and two healthy controls (12 and 15 years, each assessed in 2 replicates). In each dataset, DNA methylation differences between progeroid patients and healthy controls were assessed using the same analytical pipeline (DMPs and DMRs) used for CS samples.

### RNA extraction and Real Time (RT)-qPCR

Total RNA of WT (911VI, 198VI), UVSS (UVSS1VI, UVSTA24) and CS (CS466VI, CS548VI, CS816VI, CS177VI) cells were isolated using the RNAeasy^®^ Micro kit (Cat#74004; Qiagen) and mRNA were reverse transcribed with the SuperScript VI Reverse Transcriptase (Cat#18090050; Thermo Fisher Scientific). Resulting cDNAs were treated with RNAse H (Cat#2150B; Takara Bio) and quantified by RT-qPCR using the PowerUp^™^ SYBR^™^ Green Master mix (Cat#A25742; Thermo Fisher Scientific) on a StepOne Plus RealTime PCR system (Applied Biosystems). Data were analysed by the StepOne Plus RT PCR software v2.1 (Applied Biosystems) and normalized to TATA Binding Protein (TBP) mRNA level (=ΔCT). mRNA levels of each sample were calculated using the following formula; 2^-ΔCT^, and plotted against their DNA methylation levels. Finally, the statistical test (two-tailed) and Pearson correlation were conducted using the GraphPad Prism v6.0 software (GraphPad software).

## Acknowledgments

This work was supported by Agence Nationale de la Recherche (grant CS_AGE, aapg2019), DARRI (Institut Pasteur R&D, grant DISAGE, PasteurInnov2014), Programmes Transversales de Recherche, Institut Pasteur (grant PTR111-2017),

## Author contributions

MGB, CCh, and CCr, performed biostatistical analyses (MGB: Differential methylation analysis and comparison with other datasets; CCh and CCr: Functional analysis), analysed results and wrote the manuscript. CCr performed bench experiments and supervised the GW DNAm experiment. PG and CF provided advice for the biostatistics analysis and, together with AS discussed results and contributed to the manuscript. GL provided unpublished data on DNAm of HGPS fibroblasts, and SH provided the unpublished methylation clock for fibroblasts; both contributed to the manuscript. MR planned the experiments together with CCr, analysed results, and wrote the manuscript.

