## Supplementary Figures for "Epigenomic signature of the progeroid Cockayne syndrome exposes distinct and common features with physiological ageing"

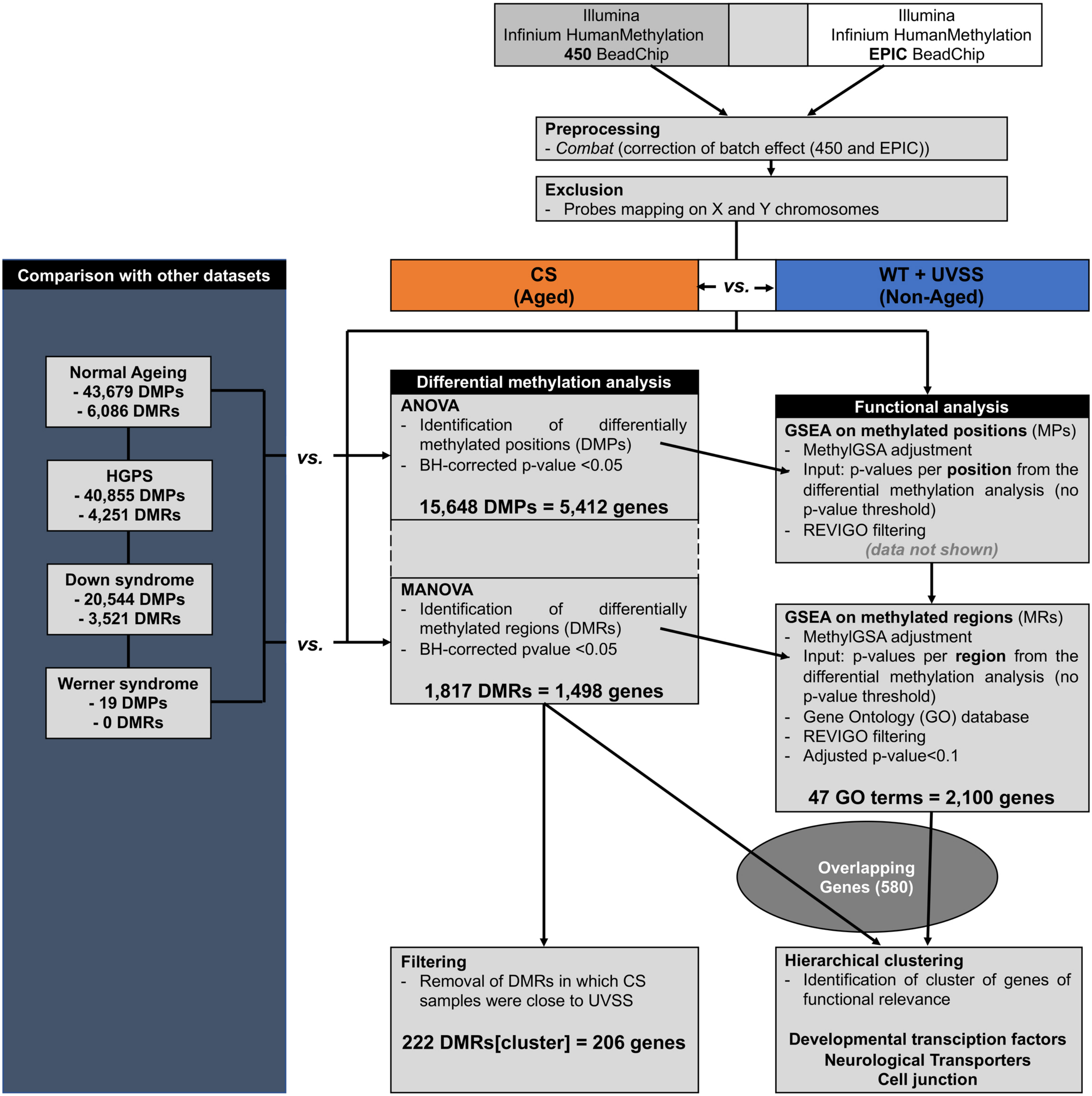


**Figure S1. Analytic pipeline for identification of the epigenomic signature of CS and common marks with physiopathological ageing**

Flow chart of the pipeline used to identify CS-associated methylation positions and regions, for the functional analysis, and for the comparison with other datasets.

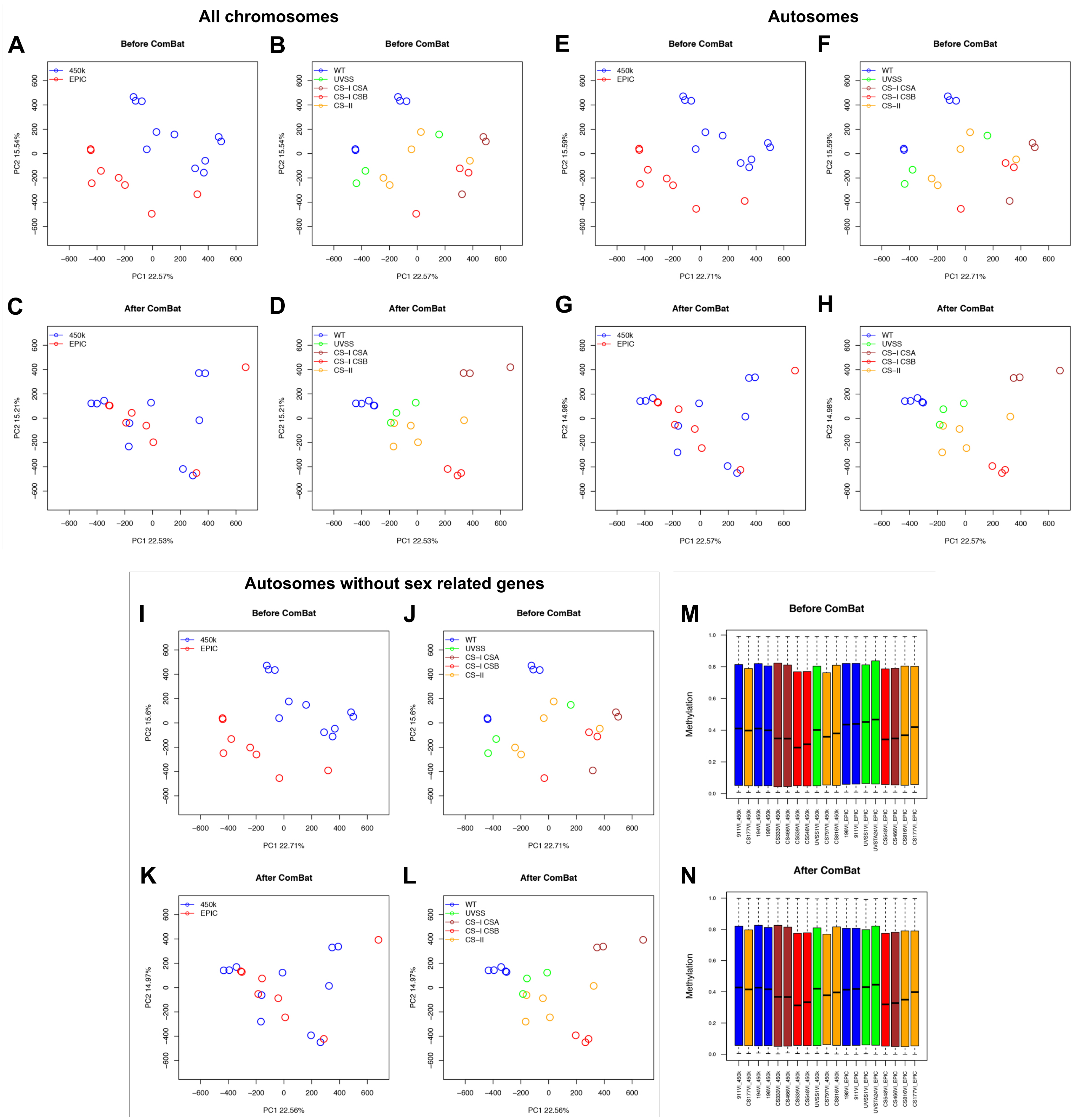


**Figure S2. Effect of sex and Combat batch correction on the global DNA methylation pattern**
PCA on methylation values of the 452567 probes common to 450k and EPIC arrays, before (**A**, **B**) and after (**C**, **D**) Combat algorithm to correct for the batch effect associated to the two platforms. In panels A and C samples are colour labelled according to microarray platform, in B and D according to the disease group. (**E-H**) Same as A-D after removing probes mapping on X and Y chromosomes. (**I-L**) Same as A-D after removing probes mapping on X and Y chromosomes and autosomic probes reported to be associated with sex. Boxplot of global DNA methylation values in each sample used for the Infinium HumanMethylation450 BeadChip and the Infinium MethylationEPIC BeadChip microarrays, before (**M**) and after (**N**) batch correction.

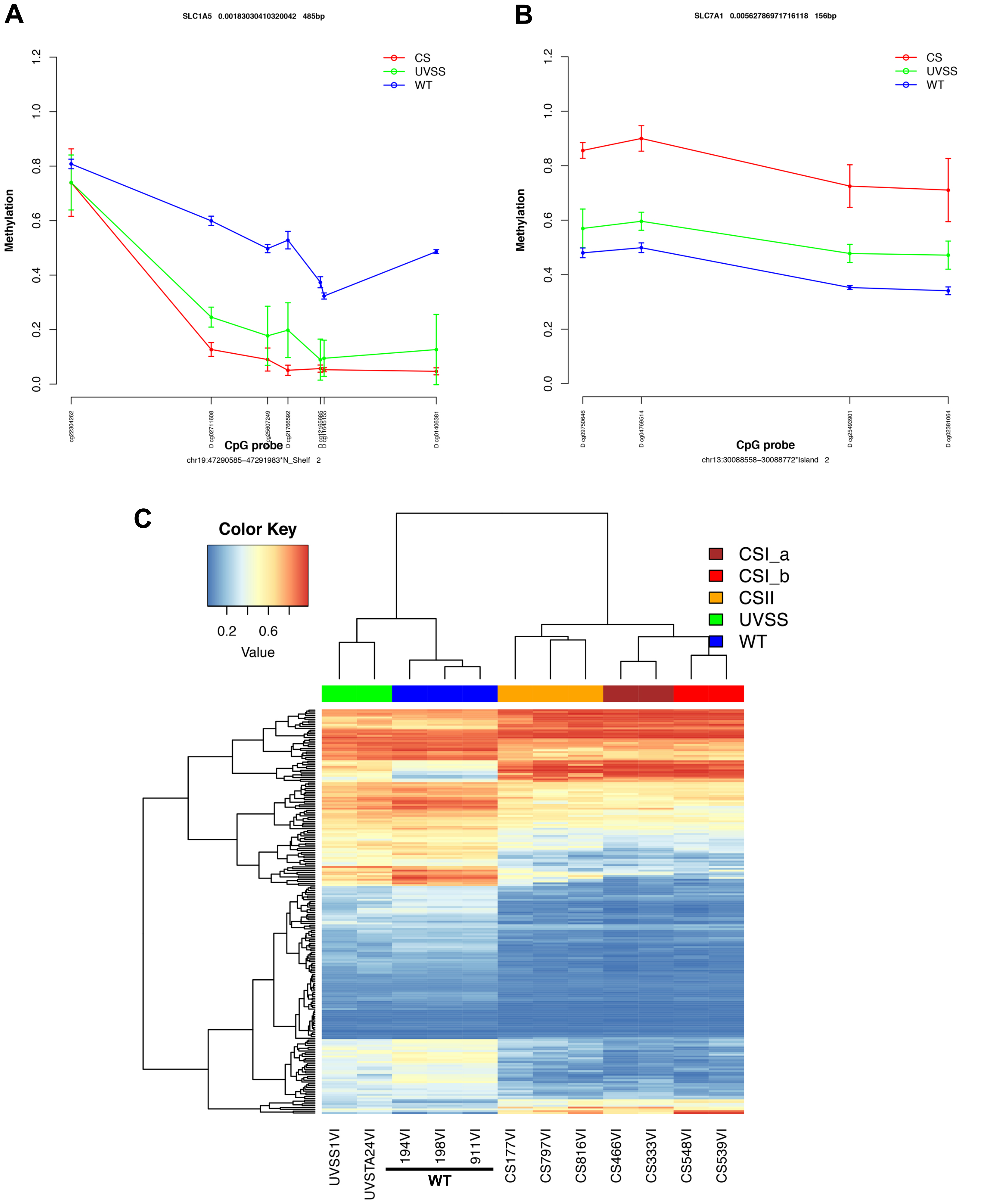


**Figure S3. Identification of stringentDMRs that unambiguously distinguished Aged *vs*. Non-Aged groups**

Lineplot displaying the methylation beta values each analysed CpG probes of WT, UVSS and CS cells in the differentially methylated region associated to (**A**) *SLC1A5* and (**B**) *SLC7A1*. (**C**) The heatmap reports DNA methylation values for the 222 stringentDMRs identified in the "Aged" versus "Non-Aged" condition (CpG probes in rows, samples in columns and color-coded). Dendrograms depicts hierarchical clustering of probes and samples.

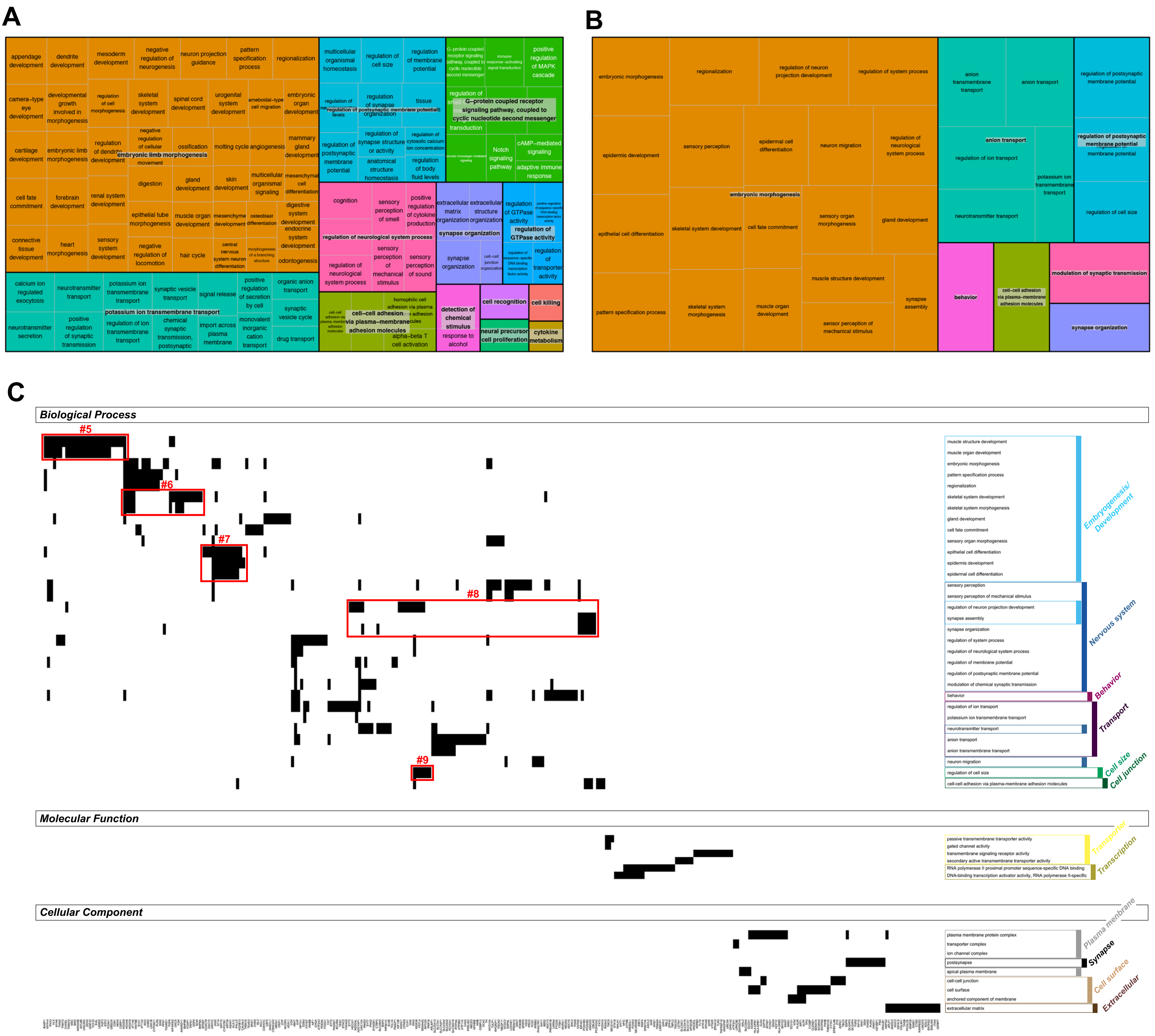


### Figure S4. Processing of the GSEA analysis

REVIGO treemap representing enriched GO terms (biological process) identified after GSEA analysis of (**A**) methylated positions and (**B**) methylated regions. Semantically similar terms are grouped in clusters identified by a "meta-term" (grey write) and by a distinct color. Rectangle areas represent the -log10 p value of each corresponding terms. (**C**) Heatmap plots reporting the relationship between the enriched GO terms identified in the Aged *vs.* Non-Aged GSEA analysis and the genes associated to DMRs belonging to these terms. Only the genes specific to each category (Process, Function, Component) are presented. The red frames indicate identified clusters of genes.
